# CD47 prevents Rac-mediated phagocytosis through Vav1 dephosphorylation

**DOI:** 10.1101/2025.02.11.637707

**Authors:** Wyatt D Miller, Abhinava K Mishra, Connor J Sheedy, Annalise Bond, Brooke M Gardner, Denise J Montell, Meghan A Morrissey

**Affiliations:** Interdisciplinary Program in Quantitative Biology, University of California, Santa Barbara, Santa Barbara CA; Molecular Cellular and Developmental Biology Department, University of California, Santa Barbara, Santa Barbara CA

**Keywords:** macrophage, phagocytosis, Fc Receptor, IgG, CD47, SIRPα

## Abstract

CD47 is expressed by viable cells to protect against phagocytosis. CD47 is recognized by SIRPα, an inhibitory receptor expressed by macrophages and other myeloid cells. Activated SIRPα recruits SHP-1 and SHP-2 phosphatases but the inhibitory signaling cascade downstream of these phosphatases is not clear. In this study, we used time lapse imaging to measure how CD47 impacts the kinetics of phagocytosis. We found that targets with IgG antibodies were primarily phagocytosed through a Rac-based reaching mechanism. Targets also containing CD47 were only phagocytosed through a less frequent Rho-based sinking mechanism. Hyperactivating Rac2 eliminated the suppressive effect of CD47, suggesting that CD47 prevents activation of Rac and reaching phagocytosis. During IgG-mediated phagocytosis, the tyrosine kinase Syk phosphorylates the GEF Vav, which then activates the GTPase Rac to drive F-actin rearrangement and target internalization. CD47 inhibited Vav1 phosphorylation without impacting Vav1 recruitment to the phagocytic synapse or Syk phosphorylation. Macrophages expressing a hyperactive Vav1 were no longer sensitive to CD47. Together this data suggests that Vav1 is a key target of the CD47 signaling pathway.

## Introduction

The immune system is regulated by a combination of activating signals on pathogenic targets and inhibitory signals that protect healthy tissue^1^. Inhibitory immune receptors are critical for tolerance in healthy tissues, and restoring homeostasis after injury and infection^2–4^. Macrophages, effectors of the innate immune system, protect the body by phagocytosing harmful targets while robustly ignoring healthy cells. CD47 is an inhibitory signal on the surface of viable cells that prevents phagocytosis^5,6^. Compared to their activating counterparts, the inhibitory signals that protect healthy cells remain poorly understood.

IgG antibodies bind bacteria, fungi or virally infected cells and trigger phagocytosis^5,7,8^. Antibody dependent phagocytosis is also an important mechanism for many cancer immunotherapies including rituximab (CD20 antibody) and trastuzumab (Her2 antibody)^9–12^. IgG is recognized by the Fc Receptor in macrophages^13^. IgG binding triggers Fc Receptor phosphorylation and recruits the effector kinase Syk. Syk activates several downstream pathways to promote F-actin reorganization and phagocytosis.

To avoid phagocytosis of healthy cells, CD47 is a potent “Don’t Eat Me” signal expressed by nearly all viable cells in mice and humans^14–16^. CD47 suppresses phagocytosis by activating the inhibitory receptor SIRPα in macrophages^17–21^. Cells lacking CD47 are rapidly cleared from healthy tissues^14,16^. Bacteria mimic CD47 to evade the innate immune system^22–24^. CD47 is also upregulated on many cancer cells, allowing these malignant cells to evade the immune system and proliferate unchecked^14,15,25^.

Modulating the CD47 signaling pathway has broad therapeutic implications. In cancer treatment, CD47 blockades have shown promising results in several phase I and phase II clinical trials, often administered in combination with a therapeutic IgG antibody that activates phagocytosis of cancer cells^6,26–30^. CD47 blockades promote both phagocytosis of cancer cells by macrophages and cross-presentation of cancer antigen by dendritic cells^15,31,32^. CD47 blockades also improve phagocytosis of diseased vascular tissue to treat atherosclerosis and clearance of viral infections^33,34^. Conversely, activating the CD47 signaling pathway has many potential therapeutic applications. Adding CD47 to transplanted cells or materials is an exciting strategy to protect these materials from macrophage phagocytosis^35–38^. Similarly, activating the CD47 receptor, SIRPα, is a potential therapy for auto-immune disorders to counteract macrophage hyperactivation^39,40^.

Despite the immense therapeutic interest in CD47 and its receptor SIRPα, very little is known about how this inhibitory signal is transduced within the macrophage. CD47 binding positions SIRPα at the phagocytic synapse between a macrophage and its target, where SIRPα’s two intracellular immune tyrosine inhibitory motifs (ITIMs) are phosphorylated by Src family kinases^41,42^. The phosphorylated ITIM motifs recruit SHP-1 and SHP-2 phosphatase, and SHP-1 is required for CD47 to inhibit phagocytosis^17–19,43–45^. The substrates of these phosphatases during phagocytosis have not been identified. Several studies have shown that SHP-1/2 phosphatases do not directly de-activate the IgG-binding Fc Receptor^17,42,46^. Downstream, this inhibitory signaling pathway prevents activation of integrin and myosin II^42,46^. However, there are many signaling events between Fc Receptor phosphorylation and the inside out activation of integrins or myosin II, leaving several potential targets for SIRPα-bound phosphatases^5^.

In this study, we clarified the downstream signaling cascade enacted by CD47, and determined how this inhibitory pathway affects phagocytosis of antibody-bound targets. We used timelapse microscopy to characterize how CD47 affected each step in the phagocytic process. For IgG opsonized targets, we observed two previously described modes of phagocytosis regulated by different GTPases. Most targets were phagocytosed via Rac-driven reaching phagocytosis and a smaller number though Rho-driven sinking phagocytosis. Targets with CD47 were only phagocytosed through the less frequent Rho-based sinking mechanism. Inhibiting Rho eliminated phagocytosis of synthetic targets and cancer cells with CD47, while inhibiting Rac had little effect. A hyperactive Rac2 mutation (Rac2^E62K^) bypassed CD47’s inhibitory effect. This demonstrates that CD47 inhibits Rac-driven reaching phagocytosis, and suggests that the direct targets of SIRPα-bound phosphatases are upstream of Rac. We found CD47 causes dephosphorylation of Vav1, the guanine nucleotide exchange factor (GEF) that activates Rac but not Rho at the phagocytic cup. CD47 did not cause any change in Syk activation or Vav1 recruitment, suggesting that Vav1 is directly regulated by SHP phosphatases. Overall, our study provides a mechanistic model for how CD47 suppresses phagocytosis.

## Results

### CD47 slows phagocytosis and increases cup retraction and failure

To study how CD47 inhibits IgG-mediated phagocytosis, we used reconstituted, cell-mimicking engulfment targets consisting of silica beads coated with fluorescent supported lipid bilayers (Fig.1A, 1B)^42,47^. To activate Fc Receptor-mediated phagocytosis, we opsonized beads with anti-biotin IgG, which binds biotinylated lipids in the bilayers. We incorporated His-tagged CD47 onto beads via attachment to Ni-NTA-conjugated lipids, so that the CD47 extracellular domain is positioned to activate SIRPα. We then incubated primary mouse bone marrow-derived macrophages (BMDMs) with IgG or IgG+CD47 beads and measured the number of internalized beads via confocal microscopy (Fig. 1A, 1B). IgG beads were engulfed approximately two times more than IgG+CD47 beads (Fig. 1B), consistent with previous studies^42,48^.

**Figure 1:**
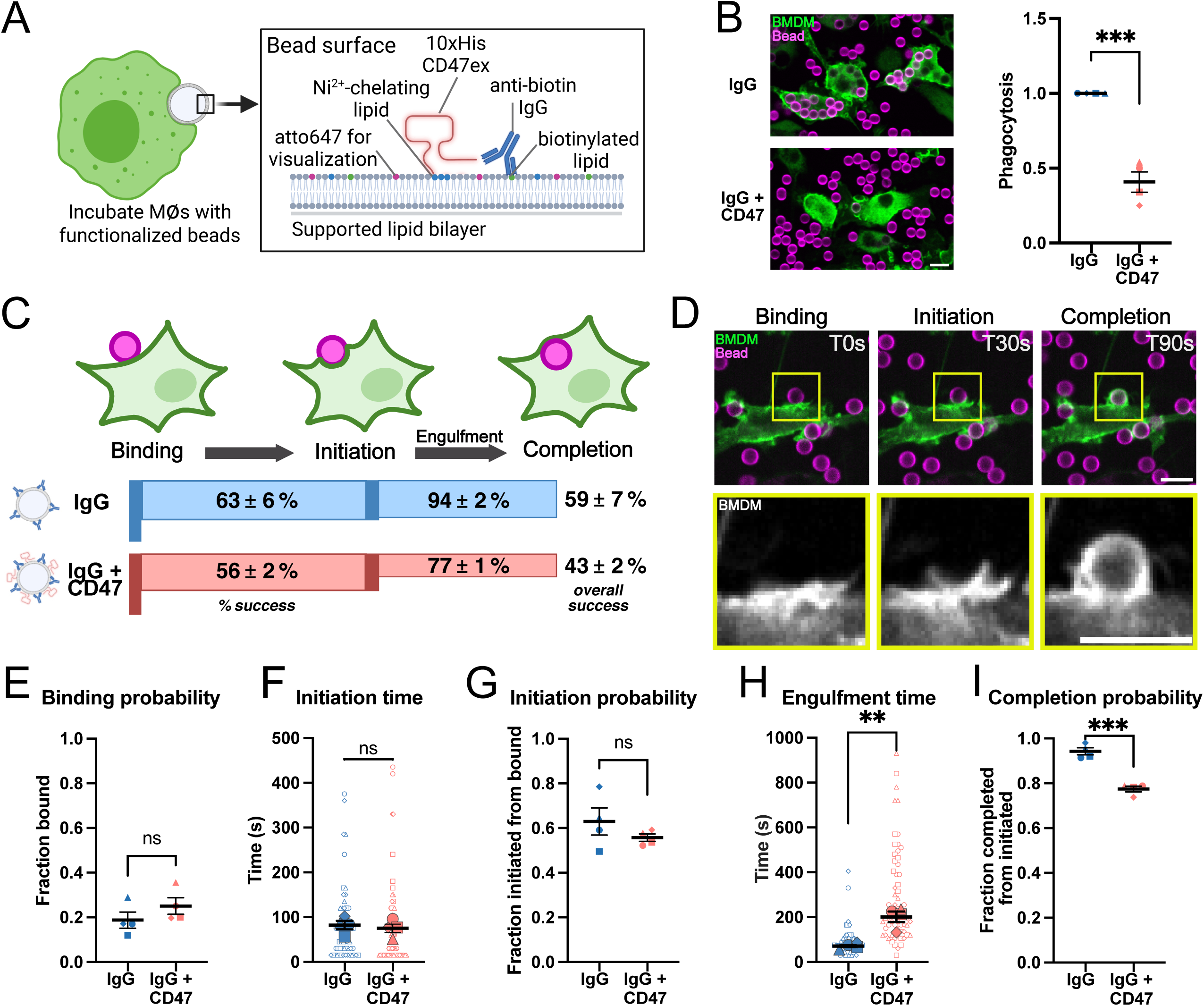
CD47 decreases the probability of completing engulfment and slows the engulfment process. (A) Schematic depicts the supported lipid bilayer system used to study phagocytosis. Silica beads are coated with a supported lipid bilayer containing fluorescent atto647 lipids. Anti-biotin IgG binds to biotinylated lipids in the bilayer. 10xHis-tagged CD47 extracellular domain is attached to beads via a Ni^2+^-chelating DGS-NTA lipid, so that the SIRPα binding domain is positioned outwards. (B) Beads (magenta) conjugated with IgG or IgG+CD47 were incubated with mouse bone marrow derived macrophages (BMDMs) expressing membrane tethered GFP (GFP-CAAX; green) for 30 minutes then imaged with confocal microscopy. The average number of beads engulfed per macrophage was counted and normalized to the maximum average number of beads engulfed per macrophage for that experiment to control for batch to batch variability in macrophage appetite. (C) Schematic depicts the stages of phagocytosis: target particle binding, initiation of phagocytosis, and completion (phagocytic cup closure). (D) Representative images of each stage. BMDMs are expressing GFP-CAAX (green, top; greyscale, bottom) and the bead supported lipid bilayer is labeled with atto647 (magenta, top). Box in the top panel shows area of inset below. Timelapse confocal microscopy was used to quantify the fraction of beads bound to a macrophage (E); the time from binding to initiation of phagocytosis, if it occurred (F); the fraction of bound beads that proceed to the initiation step (G); the engulfment time, which is the time from initiation to completion, if it completed (H); and the fraction of beads that proceeded from the initiation to completion (I). For (F) and (H), the large filled data points represent the mean of an independent replicate, while the smaller unfilled data points of the same shape indicate individual cells quantified on the same day for that replicate. For (B) each dot represents an independent replicate composed of at least 100 macrophages per condition; data was compared using an unpaired t test. For (E)-(I) each independent replicate composed of at least 15 engulfment events. The means from four independent replicates were compared using an unpaired t test. In all graphs, bars denote the mean ± SEM. * denotes p<0.05, ** denotes p<0.005, *** denotes p<0.0005. Scale bars are 10µm.

We next used time-lapse confocal microscopy to determine how CD47 affects the kinetics of phagocytosis. Phagocytosis consists of discrete steps, each associated with unique molecular mediators: (1) target particle binding, (2) initiation of phagocytosis, and (3) completion, as defined by phagocytic cup closure (Fig. 1C, D). We measured the frequency of completing each step for IgG and IgG+CD47 beads. We also measured the time between successfully completing each step (Fig. 1E-I). We found that CD47 did not inhibit target binding (Fig. 1E). We also found no difference in the frequency of initiating phagocytosis or the time between bead binding and initiation of phagocytosis (Fig. 1F, 1G). However, CD47 decreased the frequency of successfully completing phagocytosis (Fig. 1I). Most strikingly, CD47 dramatically slowed the engulfment time, or the time between initiation and completion of phagocytosis (Fig. 1H). These results indicate that CD47 affects the progression of macrophage membrane around the phagocytic target or phagocytic cup closure, rather than target particle binding or initiation of phagocytosis.

### CD47 positive targets are phagocytosed via sinking phagocytosis

Since the engulfment speed of IgG and IgG+CD47 targets was dramatically different, we hypothesized that engulfment of the two bead types may be morphologically distinct. In our timelapse data, we clearly observed two previously described forms of phagocytosis: reaching and sinking phagocytosis^49–52^. Reaching phagocytosis involves rapid extension of F-actin rich protrusions around the phagocytic target^49,52–54^. During sinking phagocytosis, the target is slowly pulled into the cell without membrane extension^49,50,52,55^. We found that IgG beads were almost always engulfed via reaching phagocytosis (Fig. 2A, C, Video S1), whereas IgG+CD47 beads were usually engulfed via sinking phagocytosis (Fig. 2B, C, Video S2). This is consistent with the slower engulfment speed of IgG+CD47 targets. To quantify the frequency of these two modes of phagocytosis more rigorously, we measured the position of the engulfment target relative to the cell cortex when phagocytosis was completed (Fig. 2D)^55^. This analysis revealed two populations of engulfed targets: one located outside the cell cortex at the time of cup closure (beads engulfed via reaching phagocytosis), and one located inside the cell cortex (beads engulfed via sinking phagocytosis). For IgG beads, 80% were outside the cell when phagocytosis completed. In contrast, the majority of IgG+CD47 targets were inside the cell when the phagocytic cup closed.

**Figure 2:**
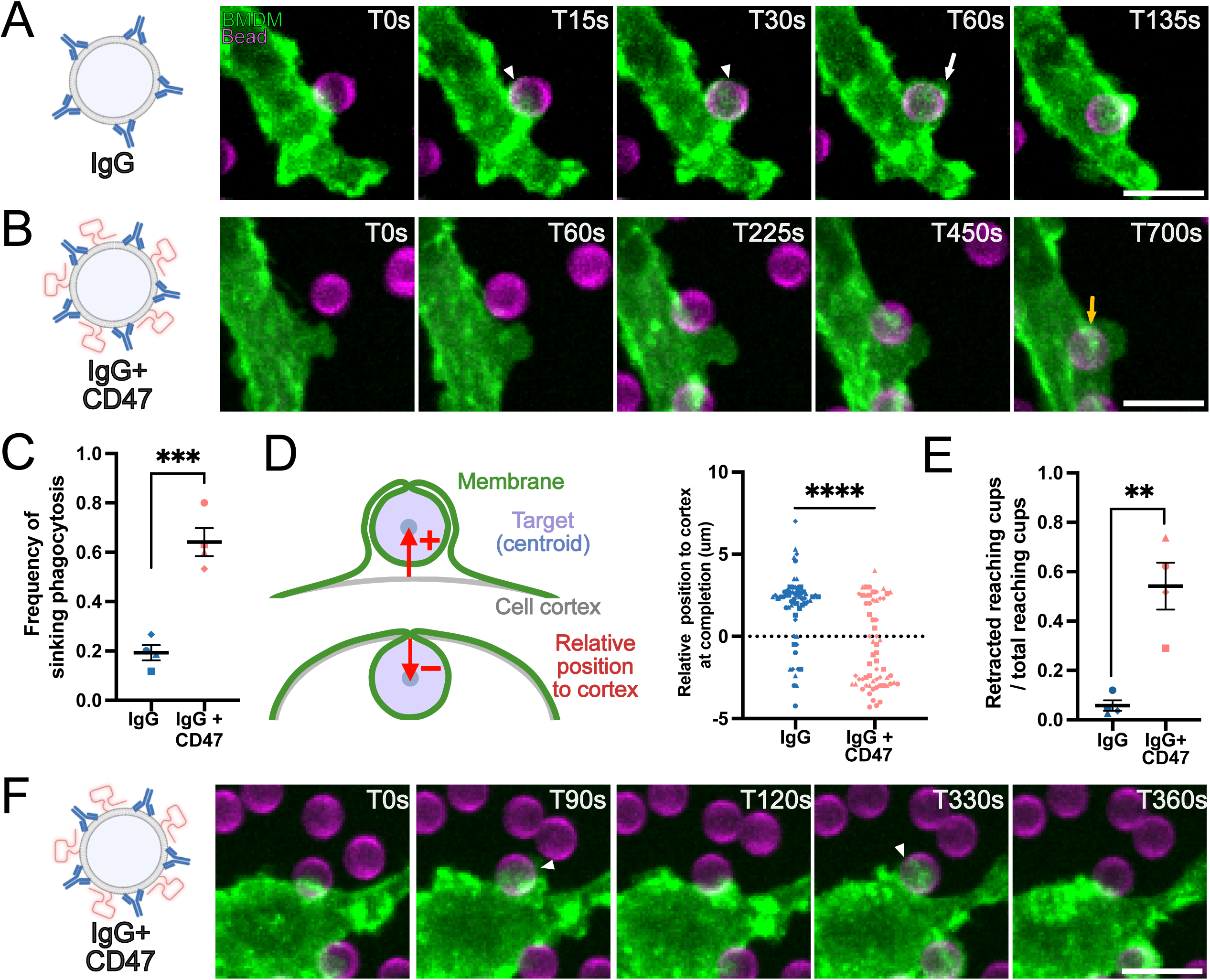
CD47 shifts primary mode of phagocytosis from reaching to sinking. (A) Timelapse images show reaching phagocytosis of an IgG bead target by a BMDM. White arrowheads point to extension of phagocyte membrane (GFP-CAAX, green) out around the target (atto647 lipid; magenta). White arrow denotes subsequent closure of the phagocytic cup while the bead remains outside the phagocyte cortex. Images correspond to Video S1 (B) Timelapse images show sinking phagocytosis of IgG+CD47 bead. Yellow arrows show closure of the phagocytic cup when the bead is within the macrophage. Images correspond to Video S2. (C) Graph depicts the fraction of successful phagocytosis that occurred via sinking phagocytosis based on the apparent morphology in timelapse confocal microscopy data. (D) Graph depicts the position of the bead centroid relative to the phagocyte cortex when phagocytosis was completed. (E) Graph depicts the fraction of retracted reaching cups out of the total number of initiated reaching cups. Retracted cups were defined as cases in our timelapse data set where membrane extensions grew around the target but were subsequently disassembled. (F) Images show an example of failed reaching phagocytosis of an IgG+CD47 bead. White arrowheads highlight extension of the macrophage membrane, which is subsequently retracted. Images correspond to Video S3. For (C) and (E), the averages from four independent experiments were plotted and compared using an unpaired t test. Bars represent the mean ± SEM. For (D), each data point represents an individual bead (*n*=60 targets per condition from 4 independent experiments), and data was compared using an unpaired t test. ** denotes p<0.005, *** denotes p<0.0005. Scale bars are 10µm.

We found that the increased failure rate of phagocytosis of CD47 targets was due to increased retraction of reaching phagocytic cups. If the macrophage initiated reaching phagocytosis of a target with CD47, the phagocytic cup usually retracted before it extended around the entire target, resulting in failed reaching phagocytosis (Fig. 2E, F; Video S3). In contrast, retraction of reaching phagocytic cups was rarely observed for IgG-only targets (Fig. 2E). Together these data indicate that CD47 inhibits reaching phagocytosis, but not sinking phagocytosis.

### Rho, not Rac, is required for phagocytosis of CD47 positive targets

Reaching and sinking phagocytosis are characterized by distinct actin cytoskeleton dynamics^52,55^. One measurable difference is that F-actin accumulates at the phagocytic cup rim during reaching but not sinking phagocytosis^55^. To test if CD47 changes F-actin organization at the phagocytic cup, we incubated BMDMs with IgG and IgG+CD47 beads, then fixed and stained with phalloidin to visualize F-actin during phagocytosis (Fig. 3A, B). Actin recruitment to phagocytic cup rims was significantly reduced during engulfment of IgG+CD47 targets compared to IgG targets (Fig. 3B). This suggests that CD47 alters actin organization during phagocytosis.

**Figure 3:**
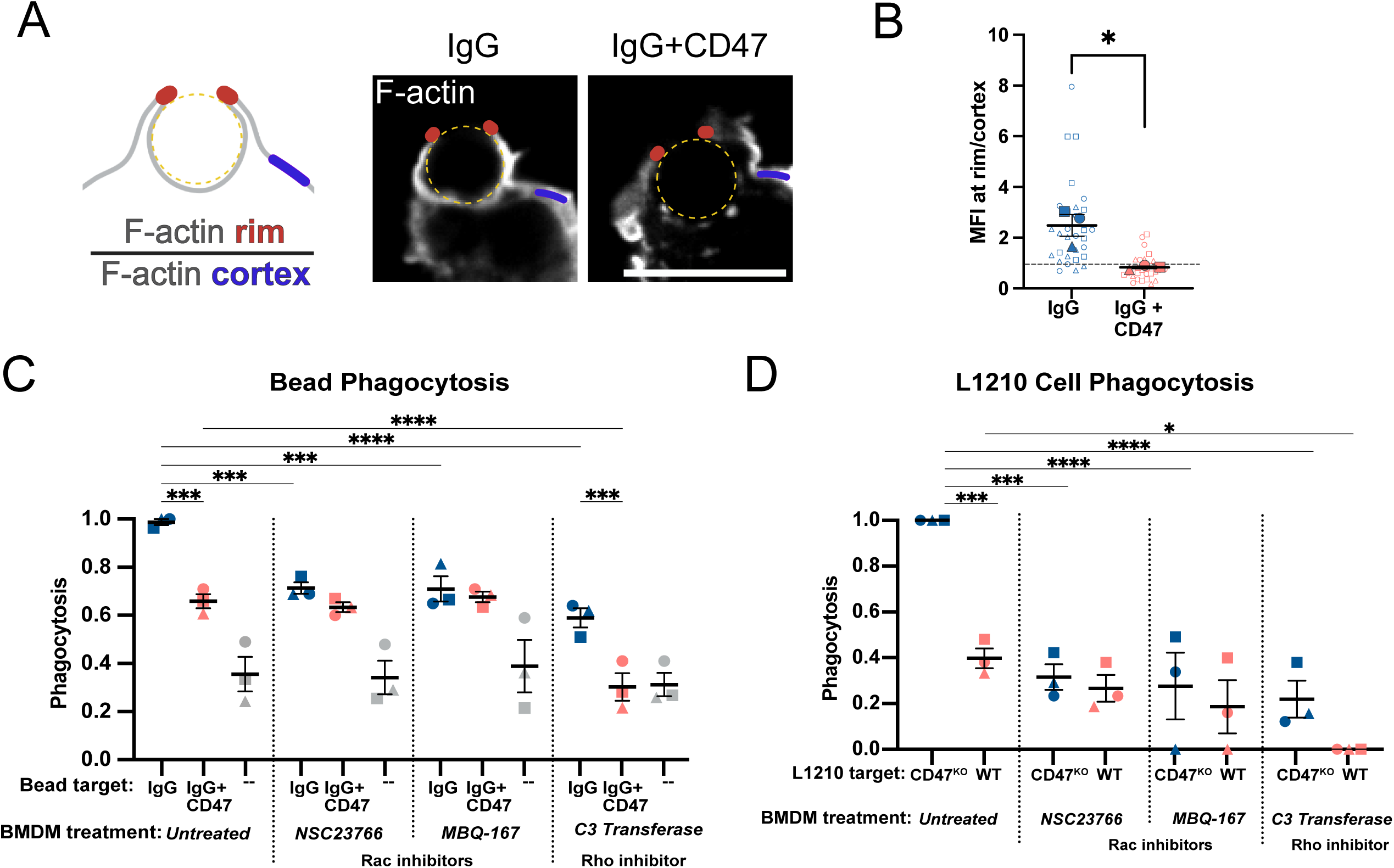
CD47 targets distinct actin regulators. (A, B) BMDMs were incubated with IgG or IgG+CD47 beads then fixed and stained with 488 phalloidin (greyscale) to visualize F-actin during phagocytosis. Enrichment of F-actin at late-stage (>50% completed) phagocytic cup rims was measured by comparing the mean fluorescent intensity (MFI) of phalloidin at the cup rim (red dot) to the MFI of phalloidin at the cell cortex (blue line). Bead position is indicated with a dashed yellow line. (B) Graph depicts data quantification described in (A). The large filled data points represent the mean of an independent replicate, while the smaller unfilled data points of the same shape indicate individual cells collected on that day for that replicate. Dashed line at 1 corresponds to no F-actin enrichment at cup rims. (C) BMDMs were treated with pharmacological inhibitors of Rac and Cdc42 (NSC23766 and MBQ-167) or Rho (C3 transferase) for 24 hours, then incubated with IgG, IgG+CD47, or unopsonized supported lipid bilayer coated beads. Phagocytosis was measured by confocal microscopy. The average number of beads phagocytosed was normalized to the maximum phagocytosis in that replicate. (D) BMDMs were treated with pharmacological inhibitors of Rac and Cdc42 (NSC23766 and MBQ-167) or Rho (C3 transferase) for 24 hours, then the inhibitors were removed and replaced with fresh media. WT or CD47^KO^ mouse L1210 leukemia cells were dyed with CellTrace Far Red then opsonized with an anti-murine CD20 monoclonal antibody and added to the BMDMs. Phagocytosis was monitored for 10 hours via timelapse microscopy. The percent of BMDMs that engulfed was quantified, and data was normalized to maximum for that experiment. For (B), the means of 4 independent experiments were compared using an unpaired t test. For (C) and (D), each data point represents an independent experiment including quantification of at least 100 macrophages. Data was compared using one-way ANOVA with Holm Sidak multiple comparison test. In all graphs, bars represent the mean ± SEM. * denotes p<0.05, ** denotes p<0.005, *** denotes p<0.0005, **** denotes p<0.00005. Scale bars are 10µm.

The Rho GTPase family (Rac, Rho and Cdc42) are regulators of the actin cytoskeleton during phagocytosis, and different GTPases have been implicated in reaching and sinking phagocytosis. Reaching phagocytosis is regulated by the GTPase Rac while sinking phagocytosis requires the GTPase Rho^52^. Since CD47 inhibits reaching phagocytosis, we hypothesized that CD47 specifically inhibits Rac, but not Rho. Additionally, since IgG+CD47 targets are engulfed via sinking phagocytosis, we hypothesized that Rho is required for phagocytosis of IgG+CD47 target particles. To test these hypotheses, we treated BMDMs with inhibitors of Rac (NSC23766 and MBQ-167; also inhibit the related GTPase CDC42) or an inhibitor of Rho (C3 transferase). We then incubated the BMDMs with IgG, IgG+CD47, or unopsonized beads (Fig. 3C). Rac inhibition significantly reduced phagocytosis of IgG targets, causing BMDMs to eat the same amount of IgG and IgG+CD47 beads. Rac inhibition did not affect phagocytosis of IgG+CD47 targets. On the other hand, Rho inhibition decreased phagocytosis of both target types, nearly eliminating phagocytosis of IgG+CD47 targets. Together, this data supports the hypothesis that CD47 inhibits Rac-mediated phagocytosis. This data also demonstrates that Rho, not Rac, is required for phagocytosis of IgG+CD47 targets.

We next sought to validate these results with cancer cell targets in lieu of synthetic bead targets. We hypothesized that CD47-expressing cancer cells, like IgG+CD47 beads, are phagocytosed through a Rho-dependent mechanism, while CD47 knockout cells, like IgG beads, are primarily phagocytosed via Rac and Cdc42. To test this, we measured phagocytosis of L1210 mouse leukemia cells opsonized with an anti-CD20 antibody. This is modeled after the clinical combination of CD20 antibodies and CD47 blockade, which has shown promise in clinical trials^27^. We used CRISPR-Cas9 to generate a monoclonal CD47^KO^ L1210 line (Supp Fig. 3). We confirmed that CD47 was not expressed on these cells by staining for CD47 (Supp Fig. 3). We again treated BMDMs with pharmacological inhibitors of Rac or Rho, then incubated them with the opsonized WT or CD47^KO^ L1210s and monitored phagocytosis via live cell microscopy (Fig. 3D). As expected, untreated BMDMs engulfed more CD47^KO^ L1210 cells than WT L1210 cells. Pharmacological inhibition of Rac decreased phagocytosis of CD47^KO^ L1210 cells, but did not affect phagocytosis of wild type L1210 cells. In contrast, Rho inhibition decreased engulfment of both CD47^KO^ and wild type L1210 cells, completely eliminating phagocytosis of wild type cells. Together, this demonstrates that CD47 inhibits Rac-mediated phagocytosis of antibody opsonized cancer cells.

### Hyperactive Rac2 overcomes CD47 inhibition

Our results show that inhibiting Rac mirrors the effect of CD47, however the direct targets of SIRPα-bound phosphatases could be upstream or downstream of Rac. We next hypothesized that if the direct targets are upstream of Rac, then hyperactivating Rac could overcome CD47-mediated inhibition of phagocytosis. In contrast, if CD47 inhibits later steps in phagocytosis, hyperactivating Rac could promote phagocytosis but targets with CD47 would still be engulfed less than cells without CD47. To test this, we isolated BMDMs from mice heterozygous for the hyperactive Rac2^E62K^ mutation (Rac2^E62K/+^)^56,57^. This Rac2 mutation is associated with a moderate increase in Rac activity^56^. We found that Rac2^E62K/+^ and wildtype macrophages phagocytosed a similar amount of antibody-opsonized CD47^KO^ L1210 cells. However, Rac2^E62K/+^ macrophages were insensitive to CD47 and phagocytosed dramatically more wild type L1210 cells compared to wild type macrophages (Fig. 4A-E; Video S4). This data demonstrates that increasing Rac activity bypasses the inhibitory effect of CD47.

**Figure 4:**
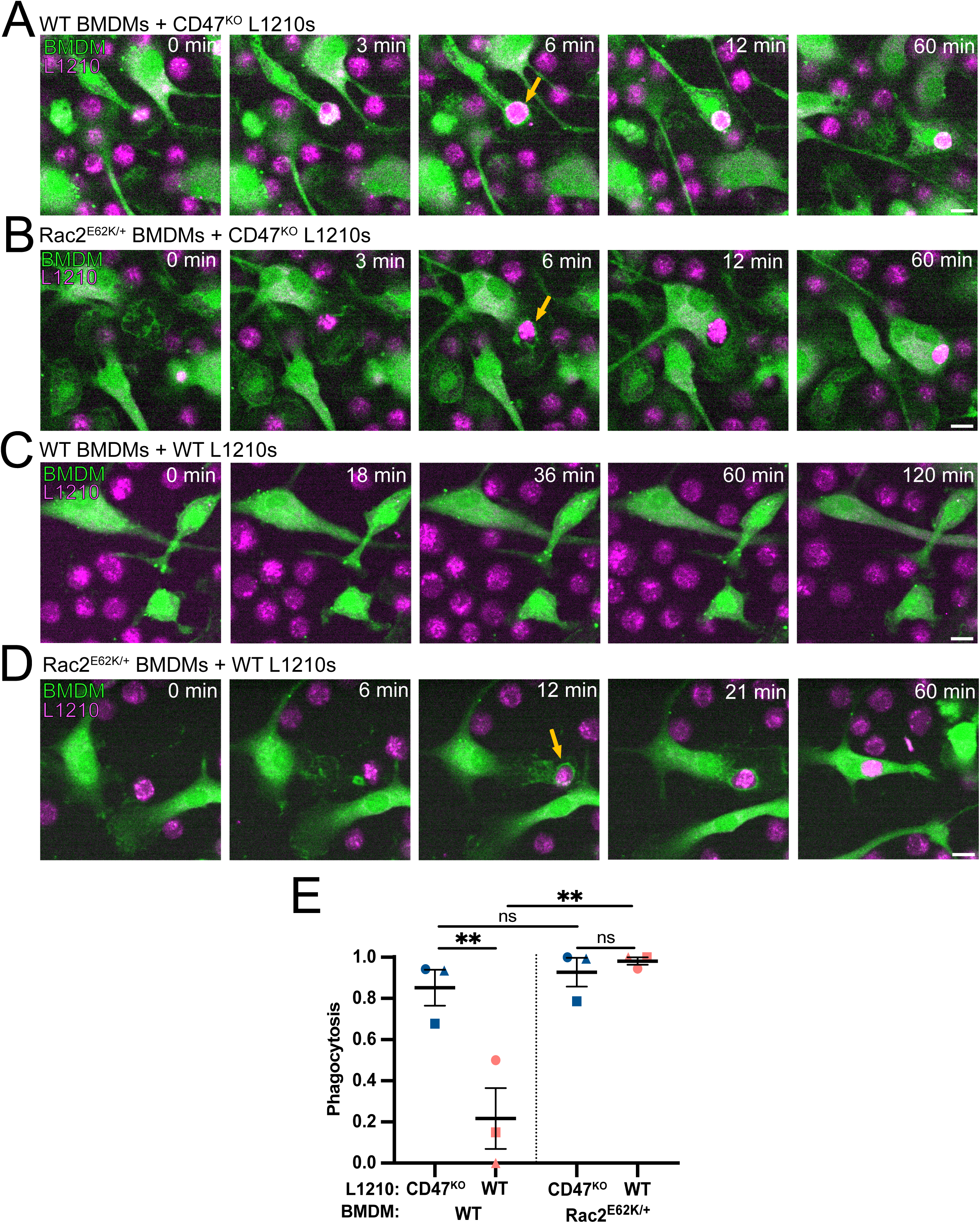
Hyperactive Rac bypasses CD47. (A-E) WT or CD47^KO^ mouse L1210 leukemia cells were dyed with CellTrace Far Red (magenta) then opsonized with an anti-murine CD20 monoclonal antibody and added to the BMDMs (green) from WT or Rac2^E62K/+^ mice. (A) Stills from a representative timelapse showing WT BMDMs incubated with CD47^KO^ L1210 cells. Yellow arrow indicates phagocytosis. (B) Stills from a representative timelapse showing Rac2^E62K/+^ BMDMs incubated with CD47^KO^ L1210 cells. Yellow arrow indicates phagocytosis. (C) Stills from a representative timelapse showing WT BMDMs incubated with CD47-positive WT L1210 cells. (D) Stills from a representative timelapse showing Rac2^E62K/+^ BMDMs incubated with CD47-positive WT L1210 cells. Yellow arrow indicates phagocytosis. (E) Graphs show the phagocytosis of L1210 cells during a 10 hour timelapse, normalized to the maximum phagocytosis on each day. Data compared using one-way ANOVA with Holm Sidak multiple comparison test. Line and bars denote mean and SEM. * denotes p<0.05, ** denotes p<0.005, *** denotes p<0.0005. Scale bars are 10µm.

### CD47 inhibits Rac through dephosphorylation of Vav1

We next considered the potential direct targets of SIRPα-bound phosphatases. We did not expect SIRPα-bound phosphatases to directly target Rac, as Rac GTPases are primarily regulated by GTP-binding and hydrolysis, rather than phosphorylation. Instead, we hypothesized that the direct target of SIRPα-bound phosphatases is upstream of Rac. Since CD47 does not inhibit Rho-mediated phagocytosis, we reasoned that the early steps in phagocytosis, required for both sinking and reaching phagocytosis, are also unlikely to be the direct target of SIRPα-bound phosphatases. After IgG binds the Fc Receptor, IgG and Fc Receptor form nanoscale clusters within the plasma membrane^58–60^. At these clusters, the Fc Receptor intracellular ITAMs are phosphorylated and recruit Syk kinase^59–61^. We have previously found that CD47 does not affect Fc Receptor clustering or alter Syk recruitment to these clusters^42^. In addition, other studies have found little or no change in Fc Receptor or Syk phosphorylation for CD47-positive targets^17,46^. To confirm this, we incubated macrophages with unopsonized, IgG, or IgG+CD47 beads, and measured Syk phosphorylation (pY346) by western blot. As expected, IgG beads increased Syk phosphorylation, and the addition of CD47 to IgG beads did not reduce the IgG-mediated Syk phosphorylation (Fig. 5A, B). Taken together, these data suggest that the direct target of SIRPα-bound phosphatases is downstream of Syk but upstream of Rac.

**Figure 5:**
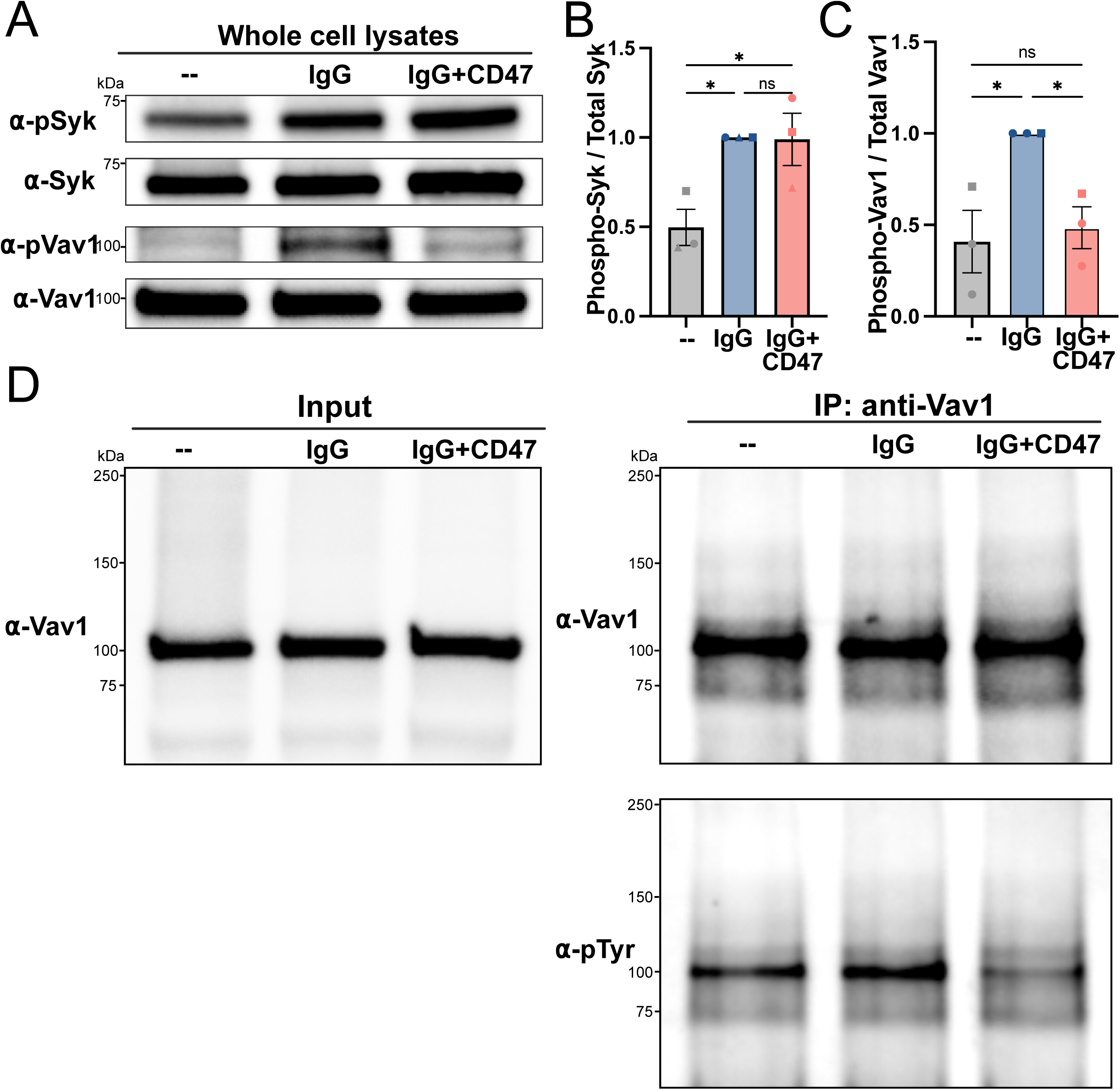
CD47 inhibits IgG-mediated phosphorylation of Vav. (A) BMDMs were incubated with unopsonized (--), IgG, or IgG+CD47 beads for 10 minutes, then lysed directly in 2X Laemmli sample buffer. Whole cell lysates were immunoblotted with the indicated antibodies. Representative of *n* = 3 independent replicates. (B, C) The ratio of phosphorylated Syk to total Syk or phosphorylated Vav1 to total Vav1 was quantified and normalized to the IgG condition for that replicate. (D) BMDMs were incubated with unopsonized (--), IgG, or IgG+CD47 beads for 10 minutes, then lysed in LB1 lysis buffer. Vav1 was immunoprecipitated, then probed for tyrosine phosphorylation. Representative of *n* = 2 independent immunoprecipitations. For (B) and (C), data was compared with one-way ANOVA with Holm Sidak multiple comparison test. Error bars denote SEM. * denotes p<0.05.

As a GTPase, Rac activation is regulated by Guanine nucleotide Exchange Factors (GEFs) that promote the exchange of GDP for GTP. Vav1 is the GEF that activates Rac during phagocytosis^62,63^. Vav1 is an attractive potential target for SIRPα-bound phosphatases for several reasons. First, Vav1 is required for Rac, but not Rho, activity during phagocytosis^64^. Second, Syk phosphorylates Vav1 on tyrosine Y174 to activate Vav1, making Vav1 the direct link between Syk and Rac^65^. Finally, in NK cells, SHP-1 directly dephosphorylates Vav1 to prevent target cell killing, suggesting that this SIRPα-bound phosphatase can directly regulate Vav1^66,67^. Supporting this, the sequence surrounding the Y174 site on Vav1 (DEIpYEDL) closely matches the optimal SHP-1 target sequences from two previous studies ((D/E)X(L/I/V)XpYXX(L/I/V) and (D/EXpY)) in both mouse and humans^68,69^. We hypothesized that SIRPα-bound phosphatases could target Vav1 to inhibit phagocytosis. We first assayed Vav1 phosphorylation using a phosphoVav1 antibody (pY174)^70,71^. We found that macrophages incubated with IgG beads showed increased Vav1 phosphorylation, and adding CD47 to the beads eliminated this increase (Fig. 5A, C). To confirm the phospho antibody staining, we immunoprecipitated Vav1 and used a pan-phosphotyrosine antibody to measure Vav1 phosphorylation (Fig. 5D, E). This confirmed that IgG increased Vav1 phosphorylation, and CD47 eliminated this increase. This data shows that CD47 eliminates IgG-induced Vav1 phosphorylation.

### CD47 does not prevent Vav1 recruitment to the phagocytic synapse

We next sought to clarify the mechanism by which CD47 reduces Vav1 phosphorylation. We began by examining Vav1 localization in the phagocytic synapse using TIRF microscopy. Vav1 binds directly to Syk, but may have other binding partners at the phagocytic cup^65,72^. In particular, Rac, which directly binds to Vav, is active throughout the actin-rich extensions that surround the phagocytic target^73–75^. We constructed planar supported lipid bilayers on a glass coverslip, then imaged IgG clustering and mCherry-Vav1. We found that on IgG bilayers, Vav1 was enriched at IgG clusters, but also present throughout the phagocytic synapse (Fig. 6A). This is consistent with the reported interaction of Vav1 with Syk, present at IgG clusters, and Rac, present along the sides of the phagocytic cup^59,73^.

**Figure 6:**
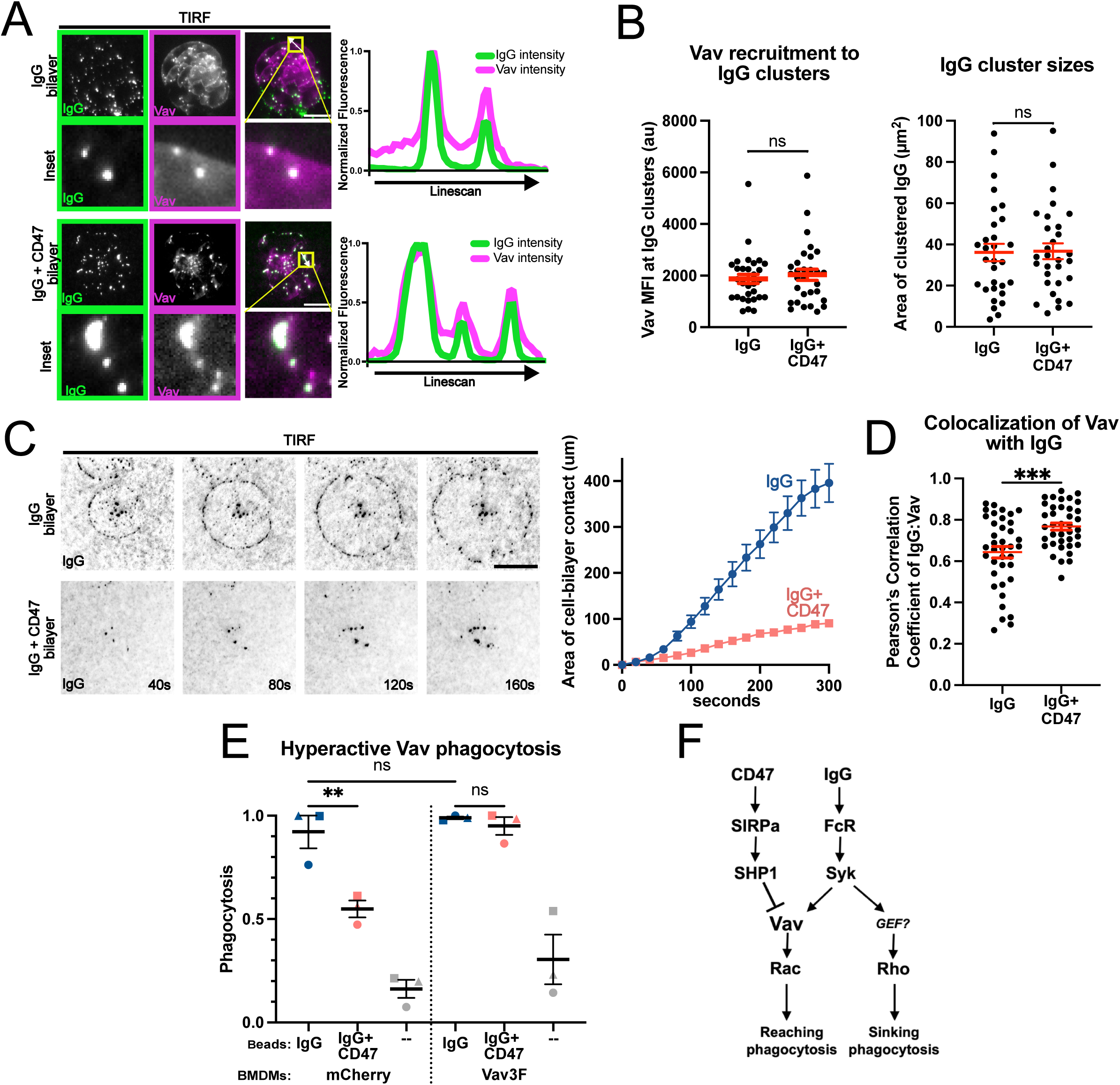
Vav1 is key target of CD47-mediated inhibition of phagocytosis. (A) TIRF images show IgG (AlexaFluor 488-IgG, green) and mCherry-Vav1 (magenta) as BMDMs interact with an IgG (top) or IgG+CD47 (bottom) bilayer. The linescan shows the fluorescent intensity of AlexaFluor 488-IgG and mCherry-Vav1 at the indicated position (white arrow in images). Intensity was normalized so that 1 is the highest observed intensity and 0 is the lowest observed intensity. (B) The mean fluorescent intensity of mCherry-Vav1 at IgG clusters (left) and the area of the IgG clusters (right) was measured for *n*=30 cells landing on IgG or IgG+CD47 bilayers, pooled from 3 independent experiments. (C) Images from TIRF microscopy timelapse show IgG (black) clustering as BMDMs land and spread on a bilayer with IgG (top) or IgG+CD47 (bottom). Graph depicts the average area of contact from *n*≥16 cells pooled from 3 separate experiments. (D) The Pearson’s Correlation Coefficient was calculated for mCherry-Vav1 and AlexaFluor 488-IgG in the footprint of *n*≥38 cells from 3 separate experiments landing on IgG or IgG+CD47 bilayers. (E) BMDMs expressing membrane-tethered mCherry (mCherry-CAAX) or a hyperactive Vav1 mutant (mCherry-Vav3F) were incubated with unopsonized (--), IgG, or IgG+CD47 beads for 30 minutes, then visualized via confocal microscopy. The average number of beads engulfed per macrophage was quantified and normalized to the highest average for that experiment. Points denote averages from each of the 3 replicates comprising at least 100 macrophages. (F) Diagram shows the proposed pathway for CD47 signaling. Data was compared using an unpaired t test (B, D) or one way ANOVA with Holm Sidak multiple comparison test. (E). Line and error bars denote mean and SEM.* denotes p<0.05, ** denotes p<0.005, *** denotes p<0.0005. Scale bars are 10µm.

SIRPα-bound phosphatases could reduce Vav1 phosphorylation either by preventing recruitment to IgG clusters where Vav1 is phosphorylated, or by dephosphorylating Vav1 directly. To distinguish between these possibilities, we used TIRF imaging to measure Vav1 recruitment to a phagocytic synapse containing CD47. We verified that CD47 inhibited cell spreading across the bilayer, demonstrating that the CD47 pathway was active (Fig. 6C). IgG cluster size was not affected by CD47, consistent with previous literature (Fig. 6A, B)^42^. CD47 did not reduce Vav1 recruitment to IgG clusters (Fig. 6A, B). In fact, Vav1 localization correlated even more strongly with IgG when CD47 was present (Fig. 6D). Since the Vav1 signal at IgG clusters is similar with IgG or IgG+CD47, this increased colocalization is likely due to a decrease in Vav1 outside of the IgG clusters. Overall, this demonstrates that CD47 does not prevent Vav1 recruitment to IgG clusters.

### Hyperactive Vav1 eliminates the effect of CD47

Since our data suggests that SHP-1 directly dephosphorylates Vav, we hypothesized that constitutive Vav1 activity would be sufficient to bypass the suppressive effect of CD47. Phosphorylation of Y174 on Vav1 displaces this residue from its intramolecular binding pocket, relieving autoinhibition of Vav1^76^. Similarly, mutating this residue to phenylalanine results in a constitutively active Vav1^77^. We used a previously characterized hyperactive Vav1 construct, where Y174 and two other key tyrosines (Y142 and Y160) are mutated to prevent Vav1 autoinhibition^77,78^. Macrophages expressing this mutant Vav1 were not sensitive to CD47, phagocytosing a comparable amount of IgG or IgG+CD47 beads (Fig. 6E). Together, this data strongly suggests that Vav1 dephosphorylation is critical for CD47 to inhibit phagocytosis.

## Discussion

CD47 is a potent “Don’t Eat Me” signal that protects healthy cells and cancer cells from macrophage attack. Understanding how CD47 inhibits phagocytosis of antibody-bound targets is particularly important, since combining CD47 blockades and monoclonal antibody therapies is the subject of several ongoing clinical trials^6,27–29^. We examined the inhibitory signaling cascade enacted by CD47. We show that CD47 specifically inhibits Rac-mediated reaching phagocytosis, but not Rho-mediated sinking phagocytosis. We consider each step in the phagocytosis signaling cascade leading to Rac activation, and demonstrate that Vav1 is the key target of SIRPα-bound phosphatases (Fig. 6F).

We show that Vav1 is dephosphorylated in response to CD47, and that constitutively active Vav1 rescues phagocytosis of CD47-positive targets (Fig. 6). This demonstrates that Vav1 is a key target of the CD47 signaling cascade. There are several reasons to think that Vav1 is directly dephosphorylated by SIRPα-bound phosphatases SHP-1 and SHP-2. After Fc Receptor activation, Syk binds to the Fc Receptor’s phosphorylated ITAMs. Vav1 binds to Syk, and is activated by Syk phosphorylation^65,72^. When CD47 is present, we find no difference in IgG cluster size, Syk recruitment or Syk phosphorylation, consistent with previous literature^17,42,46^. This suggests that the phagocytosis signaling cascade upstream of Vav1 is not affected by CD47. We further demonstrate that CD47 does not affect Vav1 recruitment to IgG clusters, arguing against an indirect effect on Vav1 due to upstream changes in signaling preventing Vav1 recruitment to the phagocytic synapse. In addition, Vav1 is directly dephosphorylated by SHP-1 in NK cells^66,67^. SHP-1 also dephosphorylates Vav1 to prevent Fas internalization and cell death in B cells^79^. We propose SHP-1 can also dephosphorylate Vav1 at the phagocytic synapse.

Our findings explain a number of results in the current literature, which could be downstream consequences of Vav1 inactivation. First, we have previously shown that CD47 prevents inside out activation of integrin^42^. Vav family GEFs are required for integrin activation during many immune cell processes including phagocytosis, cell migration and cell spreading^80–83,83–85^. Another study found that CD47 prevents myosin II phosphorylation, which is often required for phagocytosis^46^. This could be a downstream effect of Rac inhibition, since Rac promotes myosin II activation, or because myosin II is required for later steps in the phagocytosis program than Rac^86,87^. While it is possible that SIRPα-bound phosphatases target multiple proteins at the phagocytic synapse including both myosin II and Vav1, our finding that Vav1 activation rescues phagocytosis of CD47-positive targets supports the idea that Vav1 is the primary target of SIRPα-bound phosphatases.

Our work also provides some insight into Vav1 function during phagocytosis. There are very compelling studies showing that Vav1 binds directly to Syk via its SH2 domain, and this interaction promotes Vav1 phosphorylation both with purified proteins and in cells^65,72^. Vav1 phosphorylation relieves Vav1 autoinhibition^76^. Once phosphorylated via this transient interaction, Vav1 remains in the active conformation until dephosphorylated^72^. During phagocytosis, we envision that Vav1 is activated by Syk at IgG clusters, but moves on to activate Rac at other locations in the phagocytic cup. Rac is active along the sides and base of the phagocytic cup^73^. Vav1 may be targeted by SIRPα-bound phosphatases after leaving IgG clusters, since we have previously seen that SIRPα does not colocalize with IgG clusters^42^. This would explain our TIRF data showing a reduction in Vav1 at the phagocytic synapse outside of IgG clusters when CD47 is also present.

One question raised by our study is whether the mechanism of phagocytosis (reaching vs sinking phagocytosis) affects phagosome processing. Other studies have suggested that the mode of target entry can affect what proteins are recruited to the phagosome^52,88,89^. Rac and Vav1 also have known roles in phagosome processing^64^. Interestingly, CD47 blockades enhance cross presentation, which could be due to a change in phagocytosis by dendritic cells, a change in the subsequent processing of the phagocytosed material to promote cross presentation, or a separate signaling pathway in dendritic cells^32,90,91^.

Our work has broader implications for understanding inhibitory immune receptors. In macrophages, there are several other inhibitory immune receptors that regulate phagocytosis^92–96^. Future studies will need to examine whether these function through a shared mechanism, or whether each inhibitory signaling pathway has unique targets. An interesting implication of our work is that CD47 inhibits some, but not all, parts of the Fc Receptor signaling pathway. Inhibitory receptors like SIRPα are often described as the counterbalance to activating receptors like the Fc Receptors. This gives the impression of a single dial that can be turned up or turned down. Instead, we see that CD47 strongly inhibits Rac, but has no effect on Rho. This suggests a potentially more complicated situation where inhibitory receptors target specific downstream signals to sculpt the macrophage response.

Our work has several therapeutic implications. Our finding that CD47 does not inhibit Rho-mediated phagocytosis suggests that inhibitors of Rho-mediated phagocytosis would synergize with CD47. In the cases where CD47 is currently used to protect transplanted cells or particles, a secondary signal that inhibits Rho could be very potent. This also suggests that CD47 would be more protective for targets that are phagocytosed via a Rac-based mechanism than targets that are phagocytosed via a Rho-based mechanism. Cancer cells can express a variety of different “Eat Me” signals, and the phagocytic receptors that regulate engulfment can vary for different cancer populations^5^. Our work suggests that cancer cell CD47 could potently regulate phagocytosis if the signals on the cancer cell activate Rac. Finally, this work also suggests that macrophages with hyperactive Rac are insensitive to CD47, and adept at phagocytosing cancer cells^57^.

## Supplemental figure legends

**Video S1: Reaching phagocytosis of an IgG bead, related to Figure 2**. Representative video of a BMDM (green) engulfing an IgG bead (magenta) via reaching phagocytosis. Images are maximum projections of a confocal z-stack taken every 15 seconds. Scale bar is 10µm.

**Video S2: Sinking phagocytosis of an IgG+CD47 bead, related to Figure 2**. Representative image of a BMDM (green) engulfing an IgG+CD47 bead (magenta) via sinking phagocytosis. Images are maximum projections of a confocal z-stack taken every 15 seconds. Scale bar is 10µm.

**Video S3: Failed reaching phagocytosis of IgG+CD47 bead, related to Figure 2**. Representative image of a BMDM (green) attempting to engulf an IgG+CD47 bead (magenta) via reaching phagocytosis. IImages are maximum projections of a confocal z-stack taken every 15 seconds. Scale bar is 10µm.

**Video S4: Rac2^E62K/+^ macrophage phagocytoses a wild type L1210 cell, related to Figure 4**.

Representative video of a BMDM (green) phagocytosing a L1210 cell (CellTrace Far Red; magenta). Images were taken every 3 minutes. Scale bar is 10µm.

**Supplementary Figure 1: Validation of Rho GTPase inhibitors, related to Figure 3**. To validate that the GTPase inhibitors used in Figure 3 had the expected effects on macrophage actin dynamics, BMDMs were incubated for 24 hours in DMEM (untreated), or DMEM supplemented with 1mM NSC23766, 500nM MBQ-167, or 0.5µg/mL C3 transferase, then fixed and stained with GFP-conjugated phalloidin. NSC23766 and MBQ-167 had similar effects on macrophage morphology, as both induced cell rounding, shrinkage, and reduction of lamellipodia and cell polarity, characteristic of Rac inhibition. On the other hand, C3 transferase induced loss of stress fibers, protrusion of dendritic extensions and collapse of the cell body, as expected with RhoA inhibition. Scale bars are 20µm.

**Supplementary Figure 2: Validation of CD47 knockout cell line, related to Figure 3** To validate knockout of CD47 in L1210s, WT and CD47^KO^ L1210s were stained with APC-conjugated anti-CD47 and isotype control antibodies. To the left, histograms with APC signal are shown. To the right, mean fluorescent intensity of APC signal in the stained populations is graphed.

**Supplementary Figure 3: Uncropped Western blots, related to Figure 5**: (A) Uncropped Syk and phospho-Syk blots used in Figure 5A. (B) Uncropped total Vav and phospho-Vav blots used in Figure 5A. (C) To validate efficacy of Vav immunoprecipitation, the following samples were collected, run on an SDS-PAGE gel and probed for total Vav: whole cell emulsion, cell pellet, cell lysate (input used for IP), flow through, first wash, and elution (containing immunoprecipitated Vav). To the right, uncropped blots used in Figure 5D are shown.

**Supplementary Figure 4: Vav3F has morphological effects consistent with Vav1 activation, related to Figure 6** Two representative BMDMs expressing Vav3F show distinct morphology, characteristic of Vav1 hyperactivation. Vav3F-expressing cells are more spread and have increased lamellipodia and ruffles, as shown previously^78^. Vav3F-expressing cells also have large numbers of macropinosomes, as expected with increased Vav-dependent macropinocytosis. Scale bars are 20µm.

## Methods

### Key Resources Table

#### Resource Availability

##### Lead Contact

Further information and requests for reagents and resources should be directed to and will be fulfilled by the Lead Contact, Meghan Morrissey

##### Materials Availability

Plasmids generated in this study have been deposited to Addgene or can be obtained from Lead Contact.

##### Data and Code Availability

Complete imaging datasets are available from the lead contact upon request. Any additional information required to reanalyze the data reported in this paper is also available from the lead contact upon request.

#### Experimental Model and Study Participant Details

##### Cell lines

L1210 cells were obtained from the UCSF cell culture facility and certified mycoplasma and pathogen free (IMPACT III test, IDEXX BioAnalytics). The cells were cultured in DMEM (GIBCO, Catalog #11965-092) supplemented with 1% Pen-Strep-Glutamine (Corning, Catalog #30-009 CI) and 10% heat-inactivated fetal bovine serum (Atlanta Biologicals, Catalog #S11150H) at 37°C. Lenti-X 293T cells (Takara Biosciences, Cat# 632180) were cultured in DMEM, 10% FBS, 1% PSG media at 37°C.

##### Bone-marrow derived macrophage cell culture

Six- to ten-week old male and female C57BL/6 mice were sacrificed by CO_2_ inhalation. Hips and femurs were dissected and bone marrow was harvested as previously described^97^. Macrophage progenitors were differentiated for seven days in RPMI-1640 (GIBCO, Catalog # 72400120) supplemented with 10% heat-inactivated fetal bovine serum, 1% Pen-Strep-Glutamine, and 20% L929-conditioned media. Macrophage differentiation was confirmed by flow cytometry identifying CD11b and F4/80 double positive cells. Differentiated BMDMs were used for experiments from days 7 to 11.

#### Method Details

##### Lentivirus production and infection

All constructs were expressed in either BMDMs or L1210s using lentiviral infection. Lentivirus was produced in Lenti-X 293T cells transfected with pMD2.g (gift from Didier Trono, Addgene plasmid #12259 containing VSV-G envelope protein), pCMV-dR8.2 (gift from Bob Weinberg, Addgene plasmid #8455)^98^, and a lentiviral backbone containing the construct of interest using lipofectamine LTX (Invitrogen, Cat #15338-100). The media was harvested 72 h post-transfection, filtered through a 0.45 μm filter (Millapore, Cat# SLHVM33RS) and concentrated using LentiX (Takara Biosciences, Cat# 631232).

For BMDMs, concentrated lentivirus was added to cells on day 4 of differentiation, and cells were analyzed between days 7-11.

##### Supported lipid bilayer assembly

###### SUV preparation

For beads, the following chloroform-suspended lipids were mixed and desiccated overnight to remove chloroform: 95.3% POPC (Avanti, Cat# 850457), 2.5% biotinyl cap PE (Avanti, Cat# 870273), 2% DGS-NTA (Avanti, Cat# 790404), 0.1% PEG5000-PE (Avanti, Cat# 880230), and either 0.1% atto390-DOPE (ATTO-TEC GmbH, Cat# AD 390-161) or 0.1% atto647-DOPE (ATTO-TEC GmbH, Cat# AD 647–151). For planar bilayers, the following chloroform-suspended lipids were mixed and desiccated overnight to remove chloroform: 97.8% POPC (Avanti, Cat# 850457), 1% biotinyl cap PE (Avanti, Cat# 870273), 1% DGS-NTA (Avanti, Cat# 790404), 0.1% PEG5000-PE (Avanti, Cat# 880230), and 0.1% atto647-DOPE (ATTO-TEC GmbH, Cat# AD 647–151). The lipid sheets were resuspended in PBS, pH7.2 (GIBCO, Cat# 20012050) at 10mM concentration and stored under inert nitrogen gas. The lipids were broken into small unilamellar vesicles via several rounds of freeze-thaws. The lipids were then stored at -80°C under nitrogen. To remove aggregated lipids, the solution was diluted to 2 mM and filtered through a 0.22 uM filter (Millapore, Cat# SLLG013SL) immediately prior to use.

###### Planar bilayer preparation for TIRF microscopy

Ibidi coverslips (Cat# 10812) were piranha cleaned. Supported lipid bilayers were assembled in custom PDMS (Dow Corning, cat# 3097366-0516 and 3097358-1004) chambers at room temperature for 1 h. Assembled bilayers were washed 6x with PBS, then blocked with 0.2% casein (Sigma, cat# C5890) in PBS for 15 minutes. Anti-biotin IgG (Jackson ImmunoResearch Laboratories Cat# 200-002-211) was added at 1 nM, and 10x-His tagged CD47 (Sino Biological, Cat# 57231-M49H-B) was added at 10 nM. Proteins were coupled to the bilayer for 45 min. Bilayers were then washed 4x with 0.2% casein in PBS, then 2x with serum-free RPMI at RT. Imaging was conducted in serum-free RPMI at RT. Bilayers were assessed for mobility by either photobleaching or monitoring the mobility of single particles.

###### Bead preparation

Silica beads with a 5.01 μm diameter (9.79% solids, Bangs Laboratories, Cat# SS05003, Lot #16595) were washed several times with PBS, mixed with 1mM SUVs in PBS and incubated at room temperature for 30 min - 2 hrs with end-over-end mixing to allow for bilayer formation. Beads were then washed with PBS to remove excess SUVs and incubated in 0.2% casein (Sigma, Cat# C5890) in PBS for 15 min before protein coupling. Anti-biotin IgG (Jackson ImmunoResearch Laboratories Cat# 200-002-211) was added at 1 nM, and 10x-His tagged CD47 (Sino Biological, Cat# 57231-M49H-B) was added at 10 nM. We estimated the amount of IgG coupling to the beads by comparing the fluorescence of AlexaFluor-647 IgG to calibrated fluorescent beads (Quantum AlexaFluor647, Bangs Lab) and measured 200-360 molecules/um^2^ ^99^. The CD47 density was selected to give a consistent, strong suppression of phagocytosis. Proteins were coupled to the bilayer for 30 min at room temperature with end-over-end mixing.

##### Phagocytosis assays

###### Bead engulfment

50,000 BMDMs were plated per well in a 96-well glass bottom MatriPlate (Brooks, Cat# MGB096-1-2-LG-L) between 24-48 h prior to the experiment. ∼8 × 10^5^ beads were added to wells and engulfment was allowed to proceed for 30 min. The cells were imaged using spinning disc microscopy (40 × 0.95 NA Plan Apo air). Internalized particles were identified by their fluorescent supported lipid bilayer, and counted in ImageJ by a blinded analyzer using Blind-Analysis-Tools-1.0 ImageJ plug in^61^. For each well, at least 100 macrophages were scored. When Rho GTPase inhibitors were used, BMDMs plated 24 h earlier were incubated with either 1mM NSC23766 (APExBIO, Cat# A1952), 500nM MBQ-167 (MedChemExpress, Cat# HY-112842), or 0.5ug/mL C3 Transferase (Cytoskeleton, Cat# CT04-A) for 24 hours before beads were added to cells.

###### L1210 engulfment

40,000 WT or Rac2^+/E62K^ BMDMs were plated per well in a 96-well glass bottom plate 24-48 hours prior to the experiment. WT or CD47^KO^ L1210 cells were dyed with CellTrace Far Red (Thermo, Cat# C34572), incubated with an anti-mouse CD20 antibody (Genentech, clone 5D2) at 5ug/mL, then added to wells at 80,000 cells per well and imaged every 3 min for 10 hours. For each well, at least 100 macrophages were scored by a blind analyzer. Phagocytic macrophages were characterized as BMDMs that engulfed 1 or more whole, viable L1210 cell targets. When Rho GTPase inhibitors were used, BMDMs plated 24 h earlier were incubated with either 1mM NSC23766 (APExBIO, Cat# A1952), 500nM MBQ-167 (MedChemExpress, Cat# HY-112842), or 0.5ug/mL C3 Transferase (Cytoskeleton, Cat# CT04-A) for 24 hours. Inhibitors were then washed out before adding L1210 cells.

###### Kinetics of engulfment

GFP-CAAX expressing BMDMs were plated as described in the bead engulfment assay 24-48 hours prior to the experiment. Using ND acquisition in Elements, 2-3 positions per well were manually selected. Approximately 4 x 10^5^ beads were added and phagocytosis was imaged at 15 s intervals with 1um z-steps for 30 min. Bead binding was determined by counting the number of beads that came into and remained in contact with the cells throughout the imaging time, and is shown as a percentage of total beads. Initiation was identified by the frame in which the process of bead internalization began, indicated by membrane deforming or extending around the bead. Engulfment completion was identified by complete internalization of the bead by the macrophage. The initiation time was quantified as the amount of time between bead contact (the first frame in which the bead contacted the macrophage) and engulfment initiation (the first frame in which bead internalization was visualized) and was only measured for beads that were completely internalized by the end of the imaging time. The engulfment time was quantified as the amount of time between engulfment initiation and engulfment completion (the first frame in which the bead has been fully internalized by the cell). In Fig. 2C, an engulfment event was classified as reaching phagocytosis if the target was engulfed via membrane protrusions that extended away from the cell body of the macrophage, out around the target. Alternatively, an engulfment event was classified as sinking phagocytosis if the target was slowly pulled into the cell, in the absence of any such membrane protrusions. Analysis was blinded using Blind-Analysis-Tools-1.0 ImageJ plug in^61^.

##### TIRF imaging

After assembling bilayers in TIRF chambers as described earlier, BMDMs were removed from their culture dish using 5% EDTA in PBS and resuspended in serum-free RPMI before being added to the TIRF chamber for imaging.

###### Quantification of IgG clusters and Vav1 recruitment

After 15 min of interacting with the bilayer, cells that had spread on the bilayer surface were selected for analysis. Otsu thresholding in ImageJ was used to select IgG clusters in an unbiased manner. This selection was used to generate an ROI that was then applied to the Vav-mCherry channel. The area of the ROI (area of IgG clusters) and the mean Vav1 intensity within that ROI were measured.

###### Pearson’s Correlation Coefficient

The region of cell-bilayer contact was manually selected in ImageJ and the Pearson’s correlation coefficient between AlexaFluor488-IgG and Vav-mCherry was measured using the JaCoP plugin^100^.

###### Quantification of cell-bilayer contact area

For 6C the area of the cell contacting the bilayer was traced in ImageJ beginning with the first frame where the cell can be detected. Only cells with mobile IgG clusters were included.

##### Measuring actin recruitment to cup rims

50,000 BMDMs were plated per well in a 96-well glass bottom MatriPlate between 24-48 h prior to the experiment. ∼8 × 10^5^ beads were added to wells and engulfment was allowed to proceed for 15 minutes. Cells were then fixed with 4% PFA for 10 minutes, permeabilized with 0.1% Triton-X, and stained with 14nM acti-stain 488 phalloidin (Cytoskeleton, Cat# PHDG1). Cups that were between 50-100% complete were analyzed.

##### Generation and validation of CD47^KO^ L1210 cell line

To generate stable L1210 CD47^KO^ lines, WT L1210s were infected with a 3rd generation lentiviral backbone encoding Cas9^101^ and sgRNAs (sequence: GATAAGCGCGATGCCATGG) targeting the 2nd exon of mouse CD47. Single cells expressing Cas9 and guide RNAs were sorted for monoclonal populations. To validate knockout of CD47, surface expression of CD47 was assessed via FACS (Fig. S1).

Single-cell sorted monoclonal populations, as well as WT L1210s, were washed 2x with FACS blocking buffer (PBS, 0.5% BSA, 2mM EDTA), incubated in blocking buffer on ice for 10 minutes, then incubated with an APC-conjugated anti-CD47 antibody (BioLegend, cat# 127514) or an APC-conjugated isotype control antibody (BioLegend, cat# 400511) at 10 ug/mL for 30 minutes on ice in the dark. Cells were then washed 2x with blocking buffer and analyzed using an Attune NxT (Invitrogen).

##### Immunoblotting

∼1.6 × 10^7^ beads were added to 1 million BMDMs in one well of a 6-well plate, then 10 minutes later cells were washed with PBS and lysed directly in culture plates with 100ul 2X Laemmli sample buffer containing 2-mercaptoethanol. Cells were scraped and collected, then boiled for 3 minutes at 95°C and sonicated for 30sec at 50% amplitude (Branson). Proteins were resolved on 4-20% SDS-PAGE gels (Bio-Rad #4561095) before being transferred to 0.45 µm LF PVDF membranes using a wet-transfer method with Towbin transfer buffer (192 mM Glycine, 25 mM Tris-base, 20% methanol). Membranes were blocked in 3% BSA (Fisher Scientific #BP1605-100) in TBST (0.1% Tween-20) for 1 hour, then probed with desired primary antibody in 3% BSA in TBST overnight at 4°C with gentle agitation. The following day, membranes were washed 3×5 minutes with TBST before probing with species-specific HRP-conjugated secondary antibodies (Bio-Rad, Cat# 1706515) dissolved in 3% BSA in TBST for 1 hour. Membranes were then washed 3×5 minutes with TBST, followed by one wash with TBS before visualization of chemiluminescence using either Pierce ECL2 Western Blotting Substrate (Thermo Scientific #PI80196) or Pierce SuperSignal West Femto Substrate (Thermo Scientific #34095) on a ChemiDoc MP imaging system (Bio-Rad). The antibodies used for immunoblotting in this study are as follows: anti-Vav1 (CST, Cat#2502S), anti-phosphoVav1 (Abcam, Cat#ab76225), anti-Syk (CST, Cat#13198), anti-phosphoSyk (CST, Cat#2717), anti-phosphoTyrosine (CST, Cat#8954).

##### Immunoprecipitation

∼8 × 10^7^ beads were added to 10-15 million BMDMs in a 10cm plate, then 10 minutes later cells were washed with PBS and lysed with buffer LB1 (50 mM HEPES pH 7.6, 100 mM NaCl, 50 mM KCl, 10 mM MgCl_2_, 0.5 % IPEGAL CA-630, 0.1% Na-deoxycholate, 2 mM Na_3_VO_4_, and 20 mM NaF) containing 1X protease inhibitor (ThermoFisher, Cat# 78430), benzonase (EMD-Millipore, Cat# 101697), and 0.5 mM EDTA directly in 10cm plates. Cells were scraped and collected. Cells were incubated on ice for 15 minutes, passed 5x through a 25G hypodermic needle to shear cells and genomic DNA, then incubated another 15 minutes on ice with mixing by inversion. The resulting lysate was then spun 15,000g x 20 minutes at 4°C and the supernatant was collected. Lysate was then pre-cleared with Protein A agarose beads (Cell Signaling Technology, Cat# 9863S), that were pre-washed twice in LB1, for 30 minutes at 4°C to reduce non-specific binding. Beads were then spun down 5000g x 1 min and supernatant collected. 0.7 ug of Vav1 antibody (CST, Cat# 2502S) was added directly to the lysate and incubated for 2 hours at 4°C with gentle inversion. Protein A agarose slurry, that was pre-washed twice in buffer LB1, was added to each lysate + antibody mix and incubated for another 2 hours at 4°C with gentle inversion. The protein A agarose was then washed four times with buffer LB1 and bound proteins were eluted with elution buffer (0.2 M glycine pH 2.5) by incubating for 10 minutes shaking at 1100 rpm at 4°C. Beads were spun 5000g x 1 min and eluates were collected. The elution was performed twice and pooled together. To neutralize the pH, Tris-Cl pH 8.5 was added to a final concentration of 100 mM, followed by addition of 2X Laemmli sample buffer with 2-mercaptoethanol to prepare samples for SDS-PAGE.

##### Microscopy and analysis

Images were acquired on a spinning disc confocal microscope (Nikon Ti-Eclipse inverted microscope with a Yokogawa spinning disk unit and an Orca Fusion BT scMos camera) equipped with a 40 × 0.95 NA air and a 100 × 1.49 NA oil immersion objective. The microscope is also equipped with a piezo Z drive and an OkoLabs stage top incubator for temperature, CO_2_ and humidity control. TIRF imaging was performed with an iLas2 ring TIRF on the same microscope base and same camera. The microscope was controlled using Nikon Elements.

#### Quantification and Statistical Analysis

Statistical analysis was performed in Prism 8 (GraphPad). The statistical test used is indicated in the relevant figure legend. Sample sizes were predetermined and indicated in the relevant figure legend. In general the analyzer was blinded during analysis using either manual renaming of the files or the Blind-Analalysis-Tools-1.0 ImageJ plug in. The details of each quantification method and blinding strategy are included in the Methods section.

## Supporting information

Key Resources

## Acknowledgments

We thank Melanie Rodriguez for providing Rac2^E62K/+^ femurs for BMDM isolation. We also thank members of the Morrissey lab for critical feedback on the manuscript. Addgene plasmid 52961 was a gift from Feng Zhang. CJS was supported by the UC Santa Barbara Chancellor’s Fellowship. This work was funded by NCI 1DP1CA300850 to DJM; NIGMS R35GM146784 to BMG; University of California Cancer Research Coordinating Committee grant C23CR5592 and NIGMS R35GM146935 to MAM.

## Author contributions

Conceptualization, W.D.M., M.A.M.; Methodology, W.D.M., A.K.M., C.J.S. M.A.M.; Validation, W.D.M.; Formal Analysis W.D.M.; Investigation, W.D.M., A.K.M., C.J.S., AB; Resources, W.D.M., B.M.G, D.J.M, M.A.M.; Writing–Original Draft, W.D.M., M.A.M.; Writing–Review and Editing, W.D.M., B.M.G, D.J.M, M.A.M.; Visualization, W.D.M.; Supervision, B.M.G, D.J.M, M.A.M.; Funding Acquisition, B.M.G, D.J.M, M.A.M.

## Supplemental Information

Document S1. Figures S1-S3

Video S1. Reaching phagocytosis of IgG bead, related to Figure 2 Video S2. Sinking phagocytosis of IgG+CD47 bead, related to Figure 2

Video S3. Failed reaching phagocytosis, related to Figure 2

Video S4: Rac2^E62K/+^ macrophage phagocytoses a wild type L1210 cell, related to Figure 4.

## References

1. Rumpret, M. et al. Functional categories of immune inhibitory receptors. Nat. Rev. Immunol. 20, 771–780 (2020).

2. Daëron, M., Jaeger, S., Du Pasquier, L. & Vivier, E. Immunoreceptor tyrosine-based inhibition motifs: a quest in the past and future. Immunol. Rev. 224, 11–43 (2008).

3. Grebinoski, S. & Vignali, D. A. A. Inhibitory receptor agonists: The future of autoimmune disease therapeutics? Curr. Opin. Immunol. 67, 1–9 (2020).

4. Schnell, A., Bod, L., Madi, A. & Kuchroo, V. K. The yin and yang of co-inhibitory receptors: toward anti-tumor immunity without autoimmunity. Cell Res. 30, 285–299 (2020).

5. Freeman, S. & Grinstein, S. Promoters and Antagonists of Phagocytosis: A Plastic and Tunable Response. Annu. Rev. Cell Dev. Biol. 37, 89–114 (2021).

6. Veillette, A. & Chen, J. SIRPα–CD47 Immune Checkpoint Blockade in Anticancer Therapy. Trends Immunol. 39, 173–184 (2018).

7. Dilillo, D. J., Tan, G. S., Palese, P. & Ravetch, J. V. Broadly neutralizing hemagglutinin stalk-specific antibodies require FcR interactions for protection against influenza virus in vivo. Nat. Med. 20, 143–151 (2014).

8. Erwig, L. P. & Gow, N. A. R. Interactions of fungal pathogens with phagocytes. Nat. Rev. Microbiol. 14, 163–176 (2016).

9. Weiskopf, K. & Weissman, I. L. Macrophages are critical effectors of antibody therapies for cancer. mAbs 7, 303–310 (2015).

10. Shi, Y. et al. Trastuzumab triggers phagocytic killing of high HER2 cancer cells in vitro and in vivo by interaction with Fcγ receptors on macrophages. J. Immunol. Baltim. Md 1950 194, 4379–4386 (2015).

11. Chen, X., Song, X., Li, K. & Zhang, T. FcγR-Binding Is an Important Functional Attribute for Immune Checkpoint Antibodies in Cancer Immunotherapy. Front. Immunol. 10, 292 (2019).

12. Uchida, J. et al. The innate mononuclear phagocyte network depletes B lymphocytes through Fc receptor-dependent mechanisms during anti-CD20 antibody immunotherapy. J. Exp. Med. 199, 1659–1669 (2004).

13. Nimmerjahn, F. & Ravetch, J. V. Fcγ receptors as regulators of immune responses. Nat. Rev. Immunol. 8, 34–47 (2008).

14. Jaiswal, S. et al. CD47 Is Upregulated on Circulating Hematopoietic Stem Cells and Leukemia Cells to Avoid Phagocytosis. Cell 138, 271–285 (2009).

15. Majeti, R. et al. CD47 Is an Adverse Prognostic Factor and Therapeutic Antibody Target on Human Acute Myeloid Leukemia Stem Cells. Cell 138, 286–299 (2009).

16. Oldenborg, P. A. et al. Role of CD47 as a marker of self on red blood cells. Science 288, 2051–4 (2000).

17. Okazawa, H. et al. Negative Regulation of Phagocytosis in Macrophages by the CD47-SHPS-1 System. J. Immunol. 174, 2004–2011 (2005).

18. Veillette, A., Thibaudeaut, E. & Latour, S. High expression of inhibitory receptor SHPS-1 and its association with protein-tyrosine phosphatase SHP-1 in macrophages. J. Biol. Chem. 273, 22719–22728 (1998).

19. Oldenborg, P.-A., Gresham, H. D. & Lindberg, F. P. Cd47-Signal Regulatory Protein α (Sirpα) Regulates Fcγ and Complement Receptor–Mediated Phagocytosis. J. Exp. Med. 193, 855–862 (2001).

20. Barclay, A. N. & Van Den Berg, T. K. The Interaction Between Signal Regulatory Protein Alpha (SIRPα) and CD47: Structure, Function, and Therapeutic Target. Annu. Rev. Immunol. 32, 25–50 (2014).

21. Jiang, P., Lagenaur, C. F. & Narayanan, V. Integrin-associated protein is a ligand for the P84 neural adhesion molecule. J. Biol. Chem. 274, 559–62 (1999).

22. Tal, M. C. et al. P66 is a bacterial mimic of CD47 that binds the anti-phagocytic receptor SIRPα and facilitates macrophage evasion by Borrelia burgdorferi. 2024.04.29.591704 Preprint at 10.1101/2024.04.29.591704 (2024).

23. Angabo, S., et al. CD47 and thrombospondin-1 contribute to immune evasion by Porphyromonas gingivalis. Proc. Natl. Acad. Sci. 121, e2405534121 (2024).

24. Sonnert, N. D. et al. A host–microbiota interactome reveals extensive transkingdom connectivity. Nature 628, 171–179 (2024).

25. Willingham, S. B., et al. The CD47-signal regulatory protein alpha (SIRPa) interaction is a therapeutic target for human solid tumors. Proc. Natl. Acad. Sci. 109, 6662–6667 (2012).

26. Ansell, S. M. et al. Phase 1 Study of the CD47 Blocker TTI-621 in Patients with Relapsed or Refractory Hematologic Malignancies. Clin. Cancer Res. clincanres. 3706.2020 (2021) doi:10.1158/1078-0432.CCR-20-3706.

27. Advani, R. et al. CD47 Blockade by Hu5F9-G4 and Rituximab in Non-Hodgkin’s Lymphoma. N. Engl. J. Med. 379, 1711–1721 (2018).

28. Lakhani, N. J. et al. Evorpacept alone and in combination with pembrolizumab or trastuzumab in patients with advanced solid tumours (ASPEN-01): a first-in-human, open-label, multicentre, phase 1 dose-escalation and dose-expansion study. Lancet Oncol. 22, 1740–1751 (2021).

29. Johnson, L. D. S. et al. Targeting CD47 in Sézary syndrome with SIRPαFc. Blood Adv. 3, 1145–1153 (2019).

30. Sikic, B. I. et al. First-in-Human, First-in-Class Phase I Trial of the Anti-CD47 Antibody Hu5F9-G4 in Patients With Advanced Cancers. J. Clin. Oncol. Off. J. Am. Soc. Clin. Oncol. 37, 946–953 (2019).

31. Tseng, D. et al. Anti-CD47 antibody-mediated phagocytosis of cancer by macrophages primes an effective antitumor T-cell response. Proc. Natl. Acad. Sci. 110, 11103–11108 (2013).

32. Liu, X. et al. CD47 blockade triggers T cell–mediated destruction of immunogenic tumors. Nat. Med. 21, 1209–1215 (2015).

33. Kojima, Y. et al. CD47-blocking antibodies restore phagocytosis and prevent atherosclerosis. Nature 536, 86–90 (2016).

34. Cham, L. B. et al. Immunotherapeutic Blockade of CD47 Inhibitory Signaling Enhances Innate and Adaptive Immune Responses to Viral Infection. CellReports 31, 107494 (2020).

35. Rodriguez, P. L. et al. Minimal ‘Self’ Peptides That Inhibit Phagocytic Clearance and Enhance Delivery of Nanoparticles. Science 339, 971–975 (2013).

36. Yamada-Hunter, S. A. et al. Engineered CD47 protects T cells for enhanced antitumour immunity. Nature 630, 457–465 (2024).

37. Stachelek, S. J. et al. The effect of CD47 modified polymer surfaces on inflammatory cell attachment and activation. Biomaterials 32, 4317–4326 (2011).

38. Sosale, N. G. et al. “Marker of Self” CD47 on lentiviral vectors decreases macrophage-mediated clearance and increases delivery to SIRPA-expressing lung carcinoma tumors. Mol. Ther. - Methods Clin. Dev. 3, 16080 (2016).

39. Sakano, Y. et al. SIRPα engagement regulates ILC2 effector function and alleviates airway hyperreactivity via modulating energy metabolism. Cell. Mol. Immunol. 21, 1158–1174 (2024).

40. Xie, M. M. et al. An agonistic anti-signal regulatory protein α antibody for chronic inflammatory diseases. Cell Rep. Med. 4, 101130 (2023).

41. Kharitonenkov, A. et al. A family of proteins that inhibit signalling through tyrosine kinase receptors. Nature 386, 181–186 (1997).

42. Morrissey, M. A., Kern, N. & Vale, R. D. CD47 ligation repositions the inhibitory receptor SIRPA to suppress integrin activation and phagocytosis. Immunity **ePub Aug** 7, 290–302.e6 (2020).

43. Fujioka, Y. et al. A novel membrane glycoprotein, SHPS-1, that binds the SH2-domain-containing protein tyrosine phosphatase SHP-2 in response to mitogens and cell adhesion. Mol. Cell. Biol. 16, 6887–99 (1996).

44. Noguchi, T. et al. Characterization of a 115-kDa protein that binds to SH-PTP2, a protein-tyrosine phosphatase with Src homology 2 domains, in Chinese hamster ovary cells. J. Biol. Chem. 271, 27652–8 (1996).

45. Myers, D. R. et al. Shp1 Loss Enhances Macrophage Effector Function and Promotes Anti-Tumor Immunity. Front. Immunol. 11, (2020).

46. Tsai, R. K. & Discher, D. E. Inhibition of ‘self’ engulfment through deactivation of myosin-II at the phagocytic synapse between human cells. J. Cell Biol. 180, 989–1003 (2008).

47. Joffe, A. M., Bakalar, M. H. & Fletcher, D. A. Macrophage phagocytosis assay with reconstituted target particles. Nat. Protoc. 2020 157 15, 2230–2246 (2020).

48. Suter, E. C. et al. Antibody:CD47 ratio regulates macrophage phagocytosis through competitive receptor phosphorylation. Cell Rep. 36, (2021).

49. Kaplan, G. Differences in the Mode of Phagocytosis with Fc and C3 Receptors in Macrophages. Scand. J. Immunol. 6, 797–807 (1977).

50. Munthe-Kaas, A. C., Kaplan, G. & Seljelid, R. On the mechanism of internalization of opsonized particles by rat Kupffer cells in vitro. Exp. Cell Res. 103, 201–212 (1976).

51. Underhill, D. M. & Goodridge, H. S. Information processing during phagocytosis. Nat. Rev. Immunol. 12, 492–502 (2012).

52. Caron, E. & Hall, A. Identification of Two Distinct Mechanisms of Phagocytosis Controlled by Different Rho GTPases. Science 282, 1717–1721 (1998).

53. Walbaum, S. et al. Complement receptor 3 mediates both sinking phagocytosis and phagocytic cup formation *via* distinct mechanisms. J. Biol. Chem. 296, 100256 (2021).

54. Jaumouillé, V., Cartagena-Rivera, A. X. & Waterman, C. M. Coupling of β2 integrins to actin by a mechanosensitive molecular clutch drives complement receptor-mediated phagocytosis. Nat. Cell Biol. 21, 1357–1369 (2019).

55. Vorselen, D. et al. Cell surface receptors TREM2, CD14 and integrin αMβ2 drive sinking engulfment in phosphatidylserine-mediated phagocytosis. 2022.07.30.502145 Preprint at 10.1101/2022.07.30.502145 (2022).

56. Hsu, A. P. et al. Dominant activating RAC2 mutation with lymphopenia, immunodeficiency, and cytoskeletal defects. Blood 133, 1977–1988 (2019).

57. Mishra, A. K. et al. Hyperactive Rac stimulates cannibalism of living target cells and enhances CAR-M-mediated cancer cell killing. Proc. Natl. Acad. Sci. U. S. A. 120, e2310221120 (2023).

58. Goodridge, H. S., Underhill, D. M. & Touret, N. Mechanisms of Fc Receptor and Dectin-1 Activation for Phagocytosis. Traffic 13, 1062–1071 (2012).

59. Lin, J. et al. TIRF imaging of Fc gamma receptor microclusters dynamics and signaling on macrophages during frustrated phagocytosis. BMC Immunol. 17, 5 (2016).

60. Sobota, A. et al. Binding of IgG-Opsonized Particles to FcγR Is an Active Stage of Phagocytosis That Involves Receptor Clustering and Phosphorylation. J. Immunol. 175, 4450–4457 (2005).

61. Kern, N., Dong, R., Douglas, S. M., Vale, R. D. & Morrissey, M. A. Tight nanoscale clustering of Fcγ receptors using DNA origami promotes phagocytosis. eLife 10, e68311 (2021).

62. Hall, A. B. et al. Requirements for Vav Guanine Nucleotide Exchange Factors and Rho GTPases in FcγR- and Complement-Mediated Phagocytosis. Immunity 24, 305–316 (2006).

63. Kiener, P. A. et al. Cross-linking of Fc gamma receptor I (Fc gamma RI) and receptor II (Fc gamma RII) on monocytic cells activates a signal transduction pathway common to both Fc receptors that involves the stimulation of p72 Syk protein tyrosine kinase. J. Biol. Chem. 268, 24442–24448 (1993).

64. Patel, J. C., Hall, A. & Caron, E. Vav Regulates Activation of Rac but Not Cdc42 during FcγR-mediated Phagocytosis. Mol. Biol. Cell 13, 1215–1226 (2002).

65. Deckert, M., Tartare-Deckert, S., Couture, C., Mustelin, T. & Altman, A. Functional and Physical Interactions of Syk Family Kinases with the Vav Proto-Oncogene Product. Immunity 5, 591–604 (1996).

66. Mesecke, S., Urlaub, D., Busch, H., Eils, R. & Watzl, C. Integration of Activating and Inhibitory Receptor Signaling by Regulated Phosphorylation of Vav1 in Immune Cells. Sci. Signal. 4, ra36–ra36 (2011).

67. Stebbins, C. C. et al. Vav1 Dephosphorylation by the Tyrosine Phosphatase SHP-1 as a Mechanism for Inhibition of Cellular Cytotoxicity. Mol. Cell. Biol. 23, 6291–6299 (2003).

68. Yang, J. et al. Structural Basis for Substrate Specificity of Protein-tyrosine Phosphatase SHP-1*. J. Biol. Chem. 275, 4066–4071 (2000).

69. Wang, P., Fu, H., Snavley, D. F., Freitas, M. A. & Pei, D. Screening Combinatorial Libraries by Mass Spectrometry. 2. Identification of Optimal Substrates of Protein Tyrosine Phosphatase SHP-1. Biochemistry 41, 6202–6210 (2002).

70. Li, Y., Hermanson, D. L., Moriarity, B. S. & Kaufman, D. S. Human iPSC-Derived Natural Killer Cells Engineered with Chimeric Antigen Receptors Enhance Anti-tumor Activity. Cell Stem Cell 23, 181–192.e5 (2018).

71. Abate, F. et al. Activating mutations and translocations in the guanine exchange factor VAV1 in peripheral T-cell lymphomas. Proc. Natl. Acad. Sci. 114, 764–769 (2017).

72. Bustelo, X. R. Vav family exchange factors: an integrated regulatory and functional view. Small GTPases 5, e973757 (2014).

73. Hoppe, A. D. & Swanson, J. A. Cdc42, Rac1, and Rac2 Display Distinct Patterns of Activation during Phagocytosis. Mol. Biol. Cell 15, 3509–3519 (2004).

74. Masters, T. A., Pontes, B., Viasnoff, V., Li, Y. & Gauthier, N. C. Plasma membrane tension orchestrates membrane trafficking, cytoskeletal remodeling, and biochemical signaling during phagocytosis. Proc. Natl. Acad. Sci. 110, 11875–11880 (2013).

75. Hornstein, I., Alcover, A. & Katzav, S. Vav proteins, masters of the world of cytoskeleton organization. Cell. Signal. 16, 1–11 (2004).

76. Aghazadeh, B., Lowry, W. E., Huang, X.-Y. & Rosen, M. K. Structural Basis for Relief of Autoinhibition of the Dbl Homology Domain of Proto-Oncogene Vav by Tyrosine Phosphorylation. Cell 102, 625–633 (2000).

77. López-Lago, M., Lee, H., Cruz, C., Movilla, N. & Bustelo, X. R. Tyrosine Phosphorylation Mediates Both Activation and Downmodulation of the Biological Activity of Vav. Mol. Cell. Biol. 20, 1678–1691 (2000).

78. Wilsbacher, J. L., Moores, S. L. & Brugge, J. S. An active form of Vav1 induces migration of mammary epithelial cells by stimulating secretion of an epidermal growth factor receptor ligand. Cell Commun. Signal. 4, 5 (2006).

79. Koncz, G., Kerekes, K., Chakrabandhu, K. & Hueber, A.-O. Regulating Vav1 phosphorylation by the SHP-1 tyrosine phosphatase is a fine-tuning mechanism for the negative regulation of DISC formation and Fas-mediated cell death signaling. Cell Death Differ. 15, 494–503 (2008).

80. Li, X. et al. The β-glucan receptor Dectin-1 activates the integrin Mac-1 in neutrophils via Vav protein signaling to promote Candida albicans clearance. Cell Host Microbe 10, 603–615 (2011).

81. Gakidis, M. A. M. et al. Vav GEFs are required for β2 integrin-dependent functions of neutrophils. J. Cell Biol. 166, 273–282 (2004).

82. Fischer, K.-D. et al. Vav is a regulator of cytoskeletal reorganization mediated by the T-cell receptor. Curr. Biol. 8, 554–S3 (1998).

83. Ardouin, L. et al. Vav1 transduces TCR signals required for LFA-1 function and cell polarization at the immunological synapse. Eur. J. Immunol. 33, 790–797 (2003).

84. García-Bernal, D. et al. Vav1 and Rac Control Chemokine-promoted T Lymphocyte Adhesion Mediated by the Integrin α4β1. Mol. Biol. Cell 16, 3223–3235 (2005).

85. Krawczyk, C. et al. Vav1 Controls Integrin Clustering and MHC/Peptide-Specific Cell Adhesion to Antigen-Presenting Cells. Immunity 16, 331–343 (2002).

86. Vorselen, D. et al. Phagocytic ‘teeth’ and myosin-II ‘jaw’ power target constriction during phagocytosis. eLife 10, e68627 (2021).

87. van Leeuwen, F. N., van Delft, S., Kain, H. E., van der Kammen, R. A. & Collard, J. G. Rac regulates phosphorylation of the myosin-II heavy chain, actinomyosin disassembly and cell spreading. Nat. Cell Biol. 1, 242–248 (1999).

88. Yu, Y., Zhang, Z., Walpole, G. F. W. & Yu, Y. Kinetics of phagosome maturation is coupled to their intracellular motility. *Commun*. Biol. 5, 1–14 (2022).

89. Zhang, Z., et al. Propulsive cell entry diverts pathogens from immune degradation by remodeling the phagocytic synapse. Proc. Natl. Acad. Sci. 120, e2306788120 (2023).

90. Yi, T. et al. Splenic Dendritic Cells Survey Red Blood Cells for Missing Self-CD47 to Trigger Adaptive Immune Responses. Immunity 43, 764–775 (2015).

91. Xu, M. M. et al. Dendritic Cells but Not Macrophages Sense Tumor Mitochondrial DNA for Cross-priming through Signal Regulatory Protein α Signaling. Immunity 47, 363–373.e5 (2017).

92. Pluvinage, J. V. et al. CD22 blockade restores homeostatic microglial phagocytosis in ageing brains. Nature 568, 187–192 (2019).

93. Gordon, S. R. et al. PD-1 expression by tumor-associated macrophages inhibits phagocytosis and tumor immunity. Nature 545, 495–499 (2017).

94. Barkal, A. A. et al. CD24 signalling through macrophage Siglec-10 is a target for cancer immunotherapy. Nature 572, 392–396 (2019).

95. Smith, B. A. H. & Bertozzi, C. R. The clinical impact of glycobiology: targeting selectins, Siglecs and mammalian glycans. Nat. Rev. Drug Discov. 20, 217–243 (2021).

96. Theruvath, J. et al. Anti-GD2 synergizes with CD47 blockade to mediate tumor eradication. Nat. Med. 28, 333–344 (2022).

97. Weischenfeldt, J. & Porse, B. Bone Marrow-Derived Macrophages (BMM): Isolation and Applications. CSH Protoc. 2008, pdb.prot5080 (2008).

98. Stewart, S. A. et al. Lentivirus-delivered stable gene silencing by RNAi in primary cells. RNA 9, 493–501 (2003).

99. Bond, A. et al. Prior Fc receptor activation primes macrophages for increased sensitivity to IgG via long-term and short-term mechanisms. Dev. Cell (2024) doi:10.1016/j.devcel.2024.07.017.

100. Bolte, S. & Cordelières, F. P. A guided tour into subcellular colocalization analysis in light microscopy. J. Microsc. 224, 213–232 (2006).

101. Sanjana, N. E., Shalem, O. & Zhang, F. Improved vectors and genome-wide libraries for CRISPR screening. Nat. Methods 11, 783 (2014).

