## Supplementary figures and images for "CD47 prevents Rac-mediated phagocytosis through Vav1 dephosphorylation"

### Supplemental Figure 1

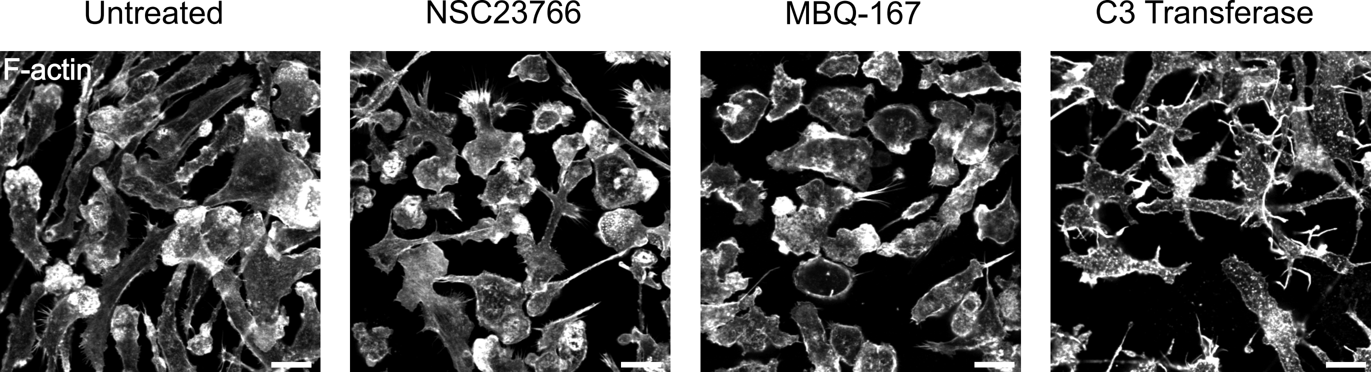

### Supplemental Figure 2

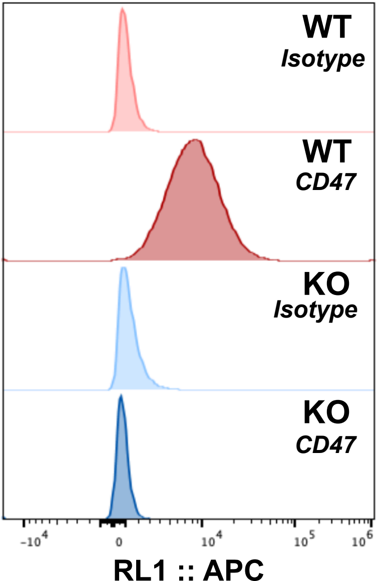

### Supplemental Figure 3

#### A Whole cell lysate - Syk blots

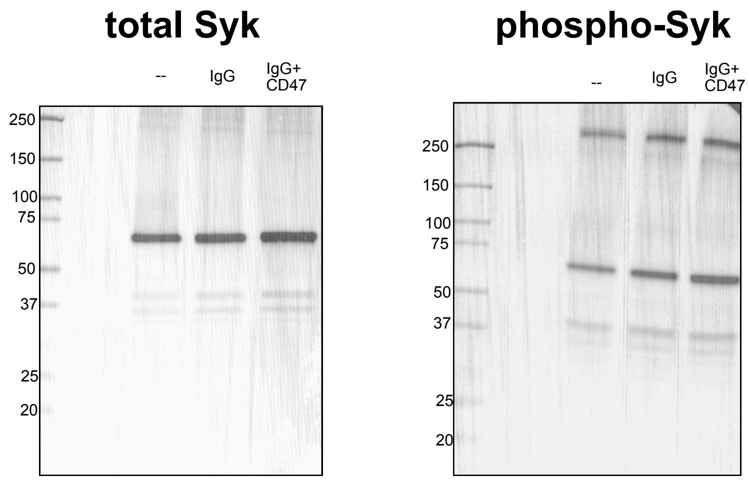


#### B Whole cell lysate - Vav1 blots

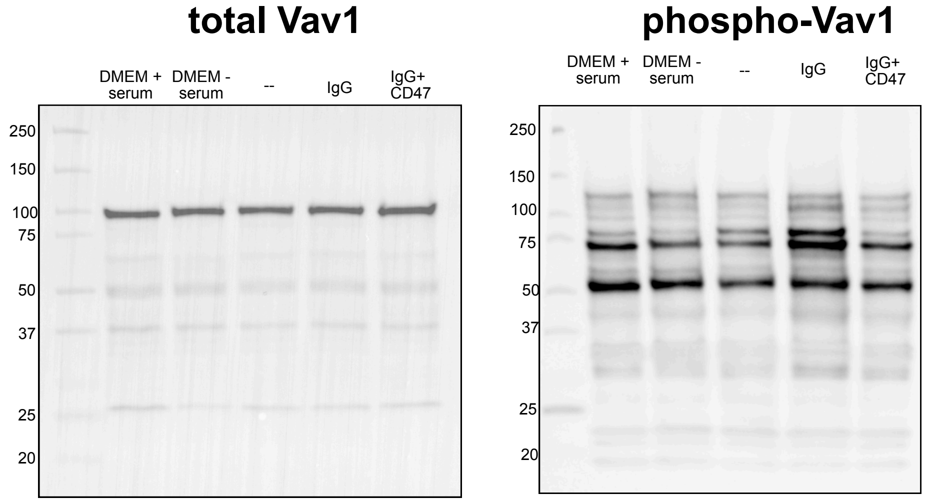


#### C Vav1 immunoprecipitation

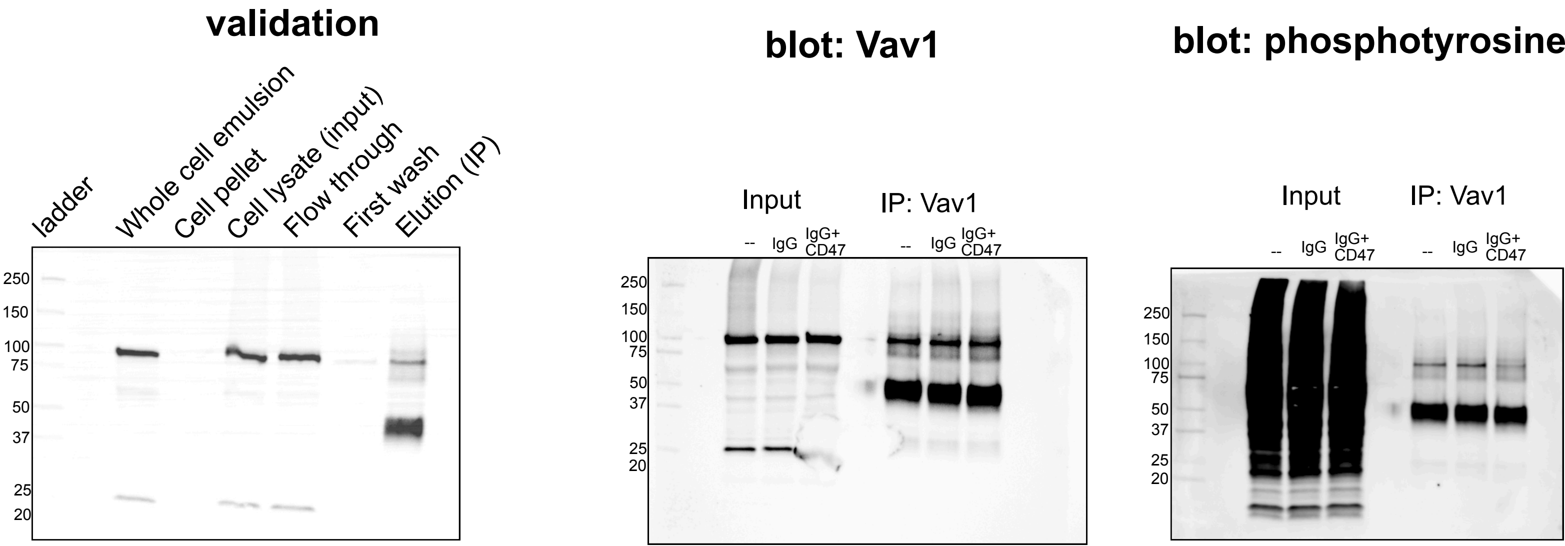

### Supplemental Figure 4

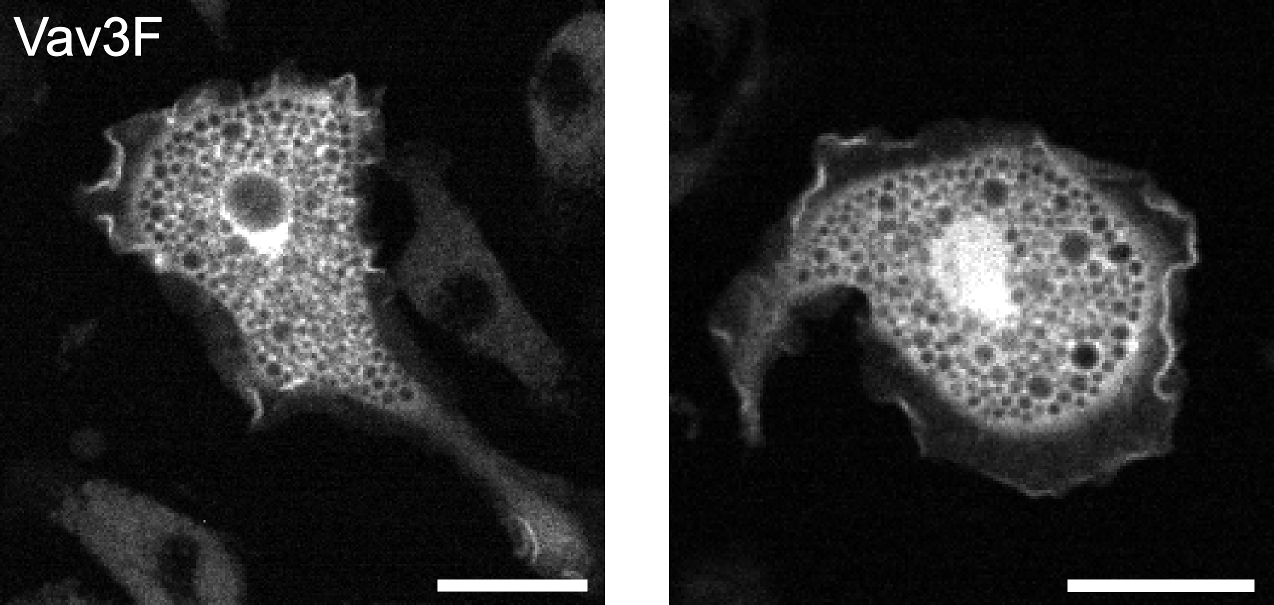
