## Supplementary material for "CD47 prevents Rac-mediated phagocytosis through Vav1 dephosphorylation": Key Resources

| REAGENT or RESOURCE | SOURCE | IDENTIFIER |
|---|---|---|
| **Antibodies** | | |
| Mouse monoclonal anti-biotin IgG | Jackson ImmunoResearch Laboratories | Cat# 200-002-211 |
| Mouse monoclonal anti-biotin IgG AlexaFluor 488 | Jackson ImmunoResearch Laboratories | Cat# 200-542-211 |
| Mouse anti-CD20 (clone 5D2) | Gift from Genentech | |
| Rat anti-F4/80 APC/Cy7 (clone BM8) | Biolegend | Cat# 123117 |
| Rabbit anti-Vav1 | Cell Signaling Technology | Cat# 2502S |
| Rabbit anti-phosphoVav1 (phospho Y174) [EP510Y] | Abcam | Cat# ab76225 |
| Rabbit anti-PhosphoTyrosine (P-Tyr-1000) | Cell Signaling Technology | Cat# 8954 |
| Rabbit anti-Syk (D3Z1E) | Cell Signaling Technology | Cat# 13198 |
| Rabbit anti-phosphoSyk (Tyr352) (clone 65E4) | Cell Signaling Technology | Cat# 2717 |
| Rat anti-CD47 APC | BioLegend | Cat# 127514 |
| Rat IgG2a, k Isotype Ctrl APC | BioLegend | Cat# 400511 |
| Goat anti-rabbit IgG (H+L)-HRP conjugate | Bio-Rad | Cat# 1706515 |
| **Chemicals, peptides, and recombinant proteins** | | |
| POPC | Avanti | Cat# 850457 |
| Biotinyl Cap PE | Avanti | Cat# 870273 |
| PEG5000-PE | Avanti | Cat# 880230 |
| Ni2+-DGS-NTA | Avanti | Cat# 790404 |
| Atto390-DOPE | ATTO-TEC GmbH | Cat# AD 390-161 |
| Atto647-DOPE | ATTO-TEC GmbH | Cat# AD 647-161 |
| CellTrace Violet | Thermo Fisher Scientific | Cat# C34571 |
| CellTrace CFSE | Thermo Fisher Scientific | Cat# C34570 |
| CellTrace Far Red | Thermo Fisher Scientific | Cat# C34572 |
| Acti-stain 488 phalloidin | Cytoskeleton, Inc | Cat# PHDG1 |
| Casein | Sigma | Cat# C5890 |
| octylphenoxypolyethoxyethanol (IPEGAL CA630) | EMD-Millipore | Cat# 8896 |
| Sodium deoxyycholate | Sigma | Cat# D6750 |
| Benzonase | EMD-Millipore | Cat# 101697 |
| Protease inhibitor | Thermo Fisher Scientific | Cat# 78430 |
| 2-mercaptoethanol | Sigma | Cat# M6250 |
| Bovine serum albumin (BSA) | Thermo Fisher Scientific | Cat# BP9703100 |
| Pierce ECL2 Western Blotting Substrate | Thermo Fisher Scientific | Cat# PI80196 |
| Pierce SuperSignal West Femto Maximum Sensitivity Substrate | Thermo Fisher Scientific | Cat# 34095 |
| Sodium fluoride | Sigma | Cat# 201154 |
| Sodium orthovanadate | Sigma | Cat# S6508-10G |
| NSC23766 trihydrochloride | APExBIO | Cat# A1952 |
| MBQ-167 | MedChemExpress | Cat# HY-112842 |
| C3 transferase | Cytoskeleton, Inc | Cat# CT04-A |
| Recombinant mouse CD47 (10xHis-tagged) | Sino Biological | Cat# 57231-M49H-B |
| **Experimental models: Cell lines** | | |
| Mouse: L1210 | ATCC | Cat# ATCC CCL-219 |
| Mouse: L1210 CD47KO | This paper | |

| | | |
|---|---|---|
| Mouse: L929 | ATCC | Cat# ATCC CLL1; RRID:CVCL_0493 |
| Human: HEK293T cells | ATCC | Cat# ATCC CRL-3216 |
| **Experimental models: Organisms/strains** | | |
| Mouse: C57BL/6 | The Jackson Laboratory | Cat# 000664 |
| Mouse: Rac2+/E62K | Provided by Amy Hsu and Steven Holland, NIAID56 | |
| **Recombinant DNA** | | |
| pHR mCherry-CAAX | Morrissey et. al.97 | mCherry fused to membrane targeting sequence from KRAB (amino acids LEKMSKDGKKKKKKSKTKCVIM) |
| pHR GFP-CAAX | This paper | GFP fused to membrane targeting sequence from KRAB (amino acids LEKMSKDGKKKKKKSKTKCVIM) |
| pMD2.G | pMD2.G was a gift from Didier Trono, Swiss Federal Institute of Technology Lausanne | Addgene plasmid # 12259; http://n2t.net/addgene:12259; RRID: Addgene_12259 |
| pCMV-dR8.2 | pCMV-dR8.2 dvpr was a gift from Bob Weinberg, Whitehead Institute for Biomedical Research | Addgene plasmid # 8455; http://n2t.net/addgene:8455; RRID: Addgene_8455 |
| pHR-mCherry-Vav1 | This paper | mCherry, Linker: GGGGSGGGGSPR, full length Vav1 ORF (Uniprot: P27870) in pHR vector |
| pHR-mCherry-Vav1-3F | This paper | As above except Y142F, Y160F, Y174F |
| lentiCRISPR-Cas9 CD47 | This paper | mouse CD47-targeting sgRNA (GATAAGCGCGATGCCATGG) cloned into lentiCRISPR v2 (Addgene: 52961109) |
| **Software and algorithms** | | |
| ImageJ-Fiji | NIH | RRID:SCR_002285; https://fiji.sc/ |
| Affinity Designer | Serif | RRID:SCR_016952 |
| Prism | Graphpad | RRID:SCR_002798 |
| Blind-Analysis-Tools-1.0 | Github | https://github.com/ahtsaJ/Blind-analysis-tools |
| JaCoP | Bolte and Cordelieres | N/A |
| FlowJo 10 | FlowJo | RRID:SCR_008520 |
| **Other** | | |
| 5.01 um silica beads | Bangs Lab | Cat# SS05003 |
| Protein A agarose beads | Cell Signaling Technology | Cat# 9863S |
| MatriPlate | Brooks | Cat# MGB09-1-2-LG-L |
| Coverslips for TIRF chambers | Ibidi | Cat# 10812 |
| PDMS | Dow Corning | Cat# 3097366-0516 |
| 4-20% stain free SDS-PAGE gels | Bio-Rad | Cat# #4568093EDU |
| 0.45 um low fluorescence PVDF membranes | Bio-Rad | cat# 1620264 |
